# Biochemical and Mechanistic Characterization of the DNA-dependent Poly(ADP-ribose) Polymerase, MoPARP1, in *Magnaporthe oryzae*

**DOI:** 10.64898/2026.09.16.752201

**Authors:** Nalleli Payne, Rachel E. Kalicharan, Adiana Reyes-Kildare, Jessie Fernandez

## Abstract

Poly(ADP-ribose) polymerases (PARPs) are important regulators of DNA repair and cellular stress responses in eukaryotes. Although mammalian PARPs have been extensively characterized, comparatively little is known about PARPs in pathogenic filamentous fungi. Here, we define biochemical features of MoPARP1 from the plant-pathogenic fungus *Magnaporthe oryzae*. Mutation of the conserved catalytic glutamate E714 abolished detectable PARylation activity, whereas PAR generated by MoPARP1 was hydrolyzed by human PARG, suggesting synthesis of polymeric ADP-ribose. Several mammalian PARP inhibitors were effective against MoPARP1 in vitro, consistent with conservation of the catalytic inhibitor-binding pocket. DNA substrates containing 5′ phosphate enhanced MoPARP1 catalytic activity, while both wild-type MoPARP1 and the catalytically inactive MoPARP1 E714A bound diverse DNA substrates. Quantitative analyses revealed that structurally distinct DNA substrates exhibit similar apparent binding affinities yet yield markedly different catalytic outputs, suggesting that DNA engagement alone does not determine productive PAR synthesis. MoPARP1 lacks the canonical N-terminal zinc-finger domains of human PARP1 and instead depends on its WGR-containing region for DNA association. Domain truncation analyses showed the BRCT-WGR region supports high-affinity DNA binding, whereas the WGR-PARP region retains catalytic competence despite weaker DNA affinity. Together, these findings establish a mechanistic framework for DNA-dependent PARylation in a filamentous fungal plant pathogen.

## Introduction

ADP-ribosylation (ADPr) is a reversible post-translational modification (PTM) with broad implications in transcription, cell signaling, chromatin organization, and DNA repair [1–3]. This modification is characterized by the enzymatic transfer of ADP-ribose units from nicotinamide adenine dinucleotide (NAD^+^) to target proteins, DNA, or RNA. The enzymes that catalyze ADP-ribose transfer are collectively known as ADP-ribosyltransferases (ARTs) [3]. Within this broad enzyme family, poly(ADP-ribose) polymerases (PARPs) represent a major ART subclass that have been extensively studied for roles in genome maintenance, transcriptional regulation, and cellular stress responses. ART family members can catalyze different forms of ADPr. In mono-ADP-ribosylation (MARylation), a single ADP-ribose unit is attached to a target molecule, whereas in poly-ADP-ribosylation (PARylation), multiple ADP-ribose units are polymerized into branched or linear chains [3]. These modifications regulate diverse cellular processes by altering protein activity, localization, stability, and interactions. Although PARPs are historically associated with PARylation, several PARP family members catalyze MARylation rather than polymer formation [4,5]. ADPr signaling is tightly regulated and rapidly reversible, in part through the activity of poly(ADP-ribose) glycohydrolases (PARGs), which hydrolyze ribose–ribose bonds within poly(ADP-ribose) chains to remove PAR modifications and terminate PAR-dependent signaling [6,7].

The ART protein family is evolutionarily widespread and has been identified in animals, bacteria, plants, and fungi. In mammals, PARPs comprise a superfamily of 17 members that can be divided into functional subclasses based on their catalytic activity, domain architecture, and cellular roles [1,8]. Among these, human PARP1, PARP2, and PARP3 (HsPARP1-3) are the core enzymes of DNA damage recognition and repair, although they differ in catalytic output: PARP1 and PARP2 are robust PARylating enzymes, whereas PARP3 primarily catalyzes MARylation [9–13]. These enzymes rapidly detect DNA damage through specialized DNA-binding regions, causing conformational changes to its catalytic region, thus initiating ADPr-dependent signaling at sites of genome damage [14]. In mammals, DNA damage-associated PARPs display distinct domain architectures. HsPARP1 contains canonical N-terminal zinc finger (ZF) DNA-binding domains together with a WGR (Trp-Gly-Arg) domain, and a C-terminal catalytic ART domain. In contrast, PARP2 and PARP3 lack ZF domains and rely more heavily on WGR-mediated DNA engagement. These architectural differences illustrate that PARP family members can use distinct DNA-binding strategies to recognize damaged DNA [15,16]. Upon binding to DNA, PARPs use NAD⁺ as a substrate, releasing nicotinamide and transferring ADP-ribose to target molecules. Within the NAD⁺-binding pocket of the ART fold, a conserved His-Tyr-Glu (HYE) catalytic triad is associated with polymerase activity. PARylation synthesis promotes the recruitment and regulation of DNA repair and chromatin-remodeling factors, orchestrating efficient DNA repair and genome integrity [15,17,18].

In plants, PARPs have been found to mediate DNA repair and responses to abiotic and biotic stress [19]. Three characterized homologs of HsPARP1-3 have been identified in *Arabidopsis thaliana*. Plant PARPs have been shown to become activated in response to oxidative damage, pathogen invasion, and environmental stressors such as drought and salinity [20]. Moreover, PARylation in plants can facilitate the recruitment of DNA repair factors and modulate hormone regulation and defense priming pathways [21].

In contrast to mammals and plants, yeast was not known to contain an active PARP ortholog or to perform PARylation under stress conditions, contributing to the assumption that yeast relies on an alternate, PARP-independent pathway for DNA repair under stress conditions [22]. However, recent work identified Pyl1, a protein with a predicted PARP catalytic domain, in *Yarrowia lipolytica* [23]. Pyl1 was shown to possess auto-PARylation activity and to PARylate the Ku70/80 complex, an essential component of the nonhomologous end-joining (NHEJ) pathway. These findings suggest that evolutionary divergent PARP-like enzymes may be present in fungi and raise important questions about the diversity of PARPs across the fungal kingdom, particularly in pathogenic fungi, where stress responses and genome integrity are critical for host colonization [24,25].

Pathogenic fungi are major contributors to crop losses, threatening the agricultural production needed to sustain a growing global population. *Magnaporthe oryzae*, the causal agent of rice blast disease, is among the most destructive fungal pathogens of rice [26,27]. Infection begins when a three-celled conidium lands on the rice surface, germinates, and differentiates a specialized infection structure known as an appressorium [28]. The appressorium generates high turgor pressure, enabling mechanical penetration of the host cuticle [28,29]. After penetration, the fungus develops invasive hyphae that proliferates inside living plant cells during the early biotrophic phase. As infection progresses, *M. oryzae* transitions to necrotrophic growth, killing host tissue and producing characteristic necrotic lesions. These lesions serve as sites for new conidium production, enabling rapid disease spread through wind- and rain-dispersed conidia. This efficient infection cycle makes rice blast difficult to control in the field, particularly because *M. oryzae* can infect nearly all above-ground tissues of the rice plant [28,29]. Severe epidemics can cause yield losses of up to 30% globally, thereby compromising rice production in regions where rice is a primary food source [30]. Management strategies, including resistant breeding, fungicides, and crop rotation, are limited in efficiency by economic costs and the fungus’ high genome plasticity [31]. Therefore, defining the molecular mechanisms that support fungal development, stress adaptation, genome maintenance, and host colonization may reveal new biological vulnerabilities for rice blast control.

Recently, a PARP homolog, MoPARP1, was identified in *M. oryzae* [32]. MoPARP1 lacks the canonical N-terminal ZF domains found in HsPARP1, suggesting that this fungal PARP uses a non-canonical DNA-binding architecture. MoPARP1 was reported to share structural similarities with HsPARPs and to exhibit PARylation activity. This work further demonstrated that MoPARP1-mediated PARylation of 14-3-3 proteins, which participate in cell signaling and gene expression regulation, is important for fungal development, appressoria formation, and activation of the MAPK pathway. More recently, biochemical characterization of PARP1 from the human pathogenic fungus *Aspergillus fumigatus* (Af-PARP1) demonstrated that fungal PARPs function as DNA-dependent enzymes whose activation is influenced by DNA architecture and 5′-terminal phosphorylation [33]. Together, these studies support an emerging role for PARylation in fungal biology and pathogenicity, and highlight the limited understanding of how DNA recognition, catalytic activation, and domain organization are coordinated among fungal PARPs. Consequently, the biochemical mechanisms underlying MoPARP1 catalytic activity, DNA binding, and domain-dependent DNA recognition remain poorly defined.

Here, we further characterize the biochemical properties of MoPARP1. We define its ADPr activity, identify catalytic residues required for activity, demonstrate that MoPARP1 binds DNA independently of catalytic activity, and determine the protein regions that contribute to DNA binding. We further examine how DNA architecture, terminal phosphorylation, and domain organization influence catalytic activation. Overall, this study reveals non-canonical features of a fungal PARP and provides new insight into the mechanisms that couple DNA recognition to productive PARylation in filamentous fungi.

## Materials and Methods

### Protein expression and purification

DNA constructs encoding MoPARP1 WT full length, E714A, BRCT-WGR, WGR, and WGR-PARP were synthesized and cloned into pET28a by Twist Bioscience (San Francisco, CA, USA). Vectors containing PARP constructs were transformed into the *E. coli* BL21 (DE3) strain. Starting cultures were grown in Lysogeny Broth (Lennox) supplemented with 50 µg/mL kanamycin at 37°C with shaking at 220 rpm until OD reached 0.8. Bacterial cultures were then supplemented with IPTG for protein induction at a final concentration of 0.4 mM and grown at 18°C for 16 hours. Bacteria was pelleted at 4°C (5,000 x g for 20 min) and stored at -80°C until further processing.

Cells were lysed with 5 mL/g of lysis buffer (50 mM sodium phosphate, pH 7.4, 300 mM NaCl, 1 mM imidazole, 1 mM DTT, and a protease inhibitor tablet (Thermo Fisher; Waltham, MA, USA) for 30 min on ice. Lysate was sonicated prior to centrifugation (20,000 × g for 30 min). Proteins of interest were then purified using nickel agarose beads. Beads were washed with 5 column volumes of wash buffer (50 mM sodium phosphate pH 7.4, 300 mM NaCl, 20 mM imidazole, 1 mM DTT) three times. Proteins were then eluted off the nickel beads using 4 mL elution buffer (50 mM sodium phosphate pH 7.4, 300 mM NaCl, 200 mM imidazole). Eluted proteins were concentrated using Prometheus centrifugal filters before loading onto ENrich 650 columns (BioRad Laboratories; Hercules, CA, USA). Proteins were stored in storage buffer (25 mM sodium phosphate pH 7.4, 50 mM NaCl, 10% glycerol) at -80°C for downstream assays.

### Bioinformatics

Protein sequences of PARP homologs from fungi, plants, and mammals were retrieved from public databases and aligned using MultAlin with default parameters [34]. Multiple sequence alignments were visualized using ESPript 3.2 [35] and subsequently edited in Adobe Illustrator 2023. Alignments were visualized and annotated to identify conserved domains, nuclear localization signals (NLSs), and catalytic residues. Predicted nuclear localization signals were identified using NLS Mapper [36]. Conserved protein domains were identified using the NCBI Conserved Domain Database (CDD) [37].

Phylogenetic relationships were inferred using the Maximum Likelihood method implemented in MEGA12 [38], and the resulting tree was visualized and edited using the Interactive Tree of Life (iTOL) platform [39]. The predicted structure of MoPARP1 was obtained from the Alphafold Structure Database (AFDB) [40]. Structural comparisons between MoPARP1 and HsPARP1 (PDB: 7KK2) were performed using the MatchMaker tool in UCSF ChimeraX with default parameters [41–43]. The resulting structural models were visualized and analyzed in UCSF ChimeraX (version 1.12; Resource for Biocomputing, Visualization, and Informatics, University of California, San Francisco, USA) [41,44].

### PARylation activity assay

PARylation assays were conducted with 100 nM PARP full-length (FL) and truncated proteins. Reactions were conducted in 50 mM Tris, pH 8, 50 mM NaCl, 1 mM DTT, 5 mM MgCl_2_, supplemented with 200 µM NAD⁺ and 200 µg/mL activated DNA (sheared salmon sperm DNA; Invitrogen, Thermo Fisher Scientific, Carlsbad, CA, USA) or 1 µM of indicated DNA oligonucleotide substrate for 30 minutes at room temperature. Reactions were stopped by the addition of 2X Laemmli SDS sample buffer and boiled for 10 minutes before resolving on 10% SDS-PAGE gels. PARylation levels were detected using anti-PARylation antibody (Cell Signaling, #89190; Danvers, MA, USA) at a 1:2000 dilution in 5% milk in TBST.

### PARP inhibitor assay

The efficacy of HsPARP inhibitors to MoPARP1 activity was assessed with a PARylation activity assay as described above. PARP inhibitors Olaparib (S1060), Veliparib (S1004), PJ34 (S7300), and Talazoparib (S7048) were purchased from Selleckchem (Houston, TX, USA), and 3-amino benzamide (sc-3501B) was purchased from Santa Cruz Biotechnology (Santa Cruz, CA, USA). Stock solutions of inhibitors were resuspended in DMSO and 0.5 µL was added to 50 µL reactions when described. Control samples contained 0.5 µL DMSO. Following immunoblot detection of PARylated proteins, PAR signal intensities were quantified by densitometric analysis using ImageJ (NIH, Bethesda, MD, USA).

### DNA substrates

DNA oligonucleotides were purchased from Eurofins Genomics (Louisville, KY, USA) (Supplementary Table S2) and resuspended in 10 mM Tris (pH 8.0), 0.1 mM EDTA, and 100 mM NaCl. Oligonucleotides were synthesized either with or without a 5′ phosphate modification, as indicated in Table S2. Selected oligonucleotides used for DNA-binding quantification were synthesized with a 3′ fluorescein (6-FAM) label. Complementary oligonucleotides were annealed by heating to 95°C for 3 min, followed by slow cooling to room temperature.

### DNA binding assay

Electromobility shift assay (EMSA) was conducted with a range of protein concentrations (1-6 µM) and a range of DNA concentrations (1-2 µM) in EMSA binding buffer (50 mM HEPES, pH 7.5, 100 mM NaCl, 0.1 mM EDTA, pH 8, 10% glycerol (v/v), 0.1 mM TCEP, 0.1 mg/mL BSA). Samples were performed in 15 µL reactions and incubated on ice for 30 min. Following incubation, tubes were spun down for 5 sec and 3 µL EMSA loading dye (10% glycerol, 10 mM Tris-HCl, pH 8, 1 mM EDTA) were added to each reaction. All samples were loaded on a 0.5% agarose gel and ran at 100 V for 30 min on ice. Gels were stained using GelRed for 5-10 min and imaged using a Chemidoc Imaging System (Bio-Rad, Hercules, CA, USA).

### NAD⁺ consumption assay

Quantification of consumed NAD⁺ was performed as previously described [45]. Briefly, reactions were conducted in 50 µL volumes using 150 nM protein in reaction buffer (50 mM Tris pH 8.0, 0.1 mg/mL BSA, 5 mM MgCl₂). The assay was performed in black propylene 96-well U-bottom microplates (Greiner Bio-One, Kremsmünster, Austria). Reactions were supplemented with 500 nM NAD⁺ and 500 nM oligonucleotides. In some cases, commercially prepared activated DNA, a DNA substrate containing strand breaks that promotes PARP activation, was used as a control at 10 µg/mL (Cytiva, Marlborough, MA, USA). Reactions were incubated in the dark with shaking at room temperature for 60 min. For NAD⁺ consumption time-course experiments, aliquots were collected at 0, 2, 5, 10, 20, 40, and 60 min. The remaining NAD⁺ was converted into a detectable signal in alkaline conditions with the addition of 20 µL 20% acetophenone in ethanol and 20 µL 2 M KOH on ice for 10 min. Finally, the samples were neutralized by adding 90 µL of formic acid and incubated on ice for 20 min before signals were read using the Biotek Synergy H1 microplate reader (BioTek Instruments, Winooski, VT, USA) at excitation and emission wavelengths of 372 nm and 444 nm, respectively. Initial NAD⁺ consumption rates (*v₀*) were determined by linear regression of the initial 10 min of the reaction (0, 2, 5, and 10 min) using GraphPad Prism and are reported as % NAD⁺ consumed min⁻¹.

### Native PAGE Electrophoretic Mobility Shift Assay (EMSA)

DNA-binding activity was assessed by native polyacrylamide gel electrophoretic mobility shift assay (PAGE-EMSA) using 3’FAM-labeled DNA substrates. Binding reactions were assembled in a final volume of 20 µL containing a fixed 5 nM concentration of DNA and increasing protein concentrations in binding buffer (10 mM HEPES, pH 7.4, 50 mM NaCl, 10% glycerol, 0.1 mM EDTA, pH 8, 1 mM DTT). The reactions were incubated on ice for 30 min to allow for complex formation. Samples were resolved on pre-run 6% native polyacrylamide gels (29:1 acrylamide:bis-acrylamide) prepared in 0.5X TBE containing 0.07% ammonium persulfate (APS) and 0.035% TEMED) at 70 V for 40 min. Gels were subsequently imaged using the fluorescein detection channel of an iBright FL1500 Imaging System (Invitrogen, Thermo Fisher Scientific, Carlsbad, CA, USA).

Integrated fluorescence intensities for free DNA and protein-DNA complexes were quantified in ImageJ/Fiji [46,47] using identical regions of interest (ROIs) across all lanes. Background fluorescence was measured from an area lacking signal and subtracted from each measurement. The fraction of DNA bound was calculated according to the previous reports [48].

Fraction-bound values were plotted as a function of protein concentration and analyzed in GraphPad Prism (version 11.0.2; GraphPad Software). Data were fitted by nonlinear regression using a one-site specific binding model to determine the apparent dissociation constant (*K_d_*). *K_d_* values are reported as the mean ± standard deviation from three independent biological experiments.

### Statistics

All experiments were independently performed three times (*n*=3 independent experiments), with three technical replicates included in each experiment. Statistical analyses were conducted using GraphPad Prism. Comparisons between two groups were performed using a two-tailed unpaired Welch’s *t*-test. For comparisons among multiple groups, data were assessed for normality using the Shapiro-Wilk test and for homogeneity of variance using the Brown-Forsythe test. When these assumptions were met, one-way ANOVA followed by Dunnett’s and Tukey’s multiple-comparisons test was performed. Data are presented as mean ± standard deviation (SD), unless otherwise indicated. Statistical significance was defined as *P* < 0.05 and indicated as follows: ns, not significant, *P* < 0.05 (\**), P < 0.01 (**), P < 0.001 (\*\*\**), and *P* < 0.0001 (****).

## Results

### MoPARP1 shares conserved domains with human PARPs

To further characterize the evolutionary and structural features of MoPARP1, we performed comparative domain, sequence, phylogenetic, and structural analyses. Previous work demonstrated that MoPARP1 possesses PARylation activity [32]. Consistent with these findings, domain analysis revealed that MoPARP1 contains conserved BRCA1 C-terminal (BRCT), tryptophan-glycine-arginine (WGR), regulatory helical (HD), and catalytic (CD) domains characteristic of DNA-dependent PARPs (Fig. 1A). Similar domain architectures were observed in human PARP1 (HsPARP1), PARP2 (HsPARP2), and PARP3 (HsPARP3). However, unlike HsPARP1, MoPARP1 lacks the N-terminal zinc-finger (ZF) domains responsible for DNA damage recognition and binding. Sequence analysis using NLS Mapper [36] identified a strong predicted monopartite nuclear localization signal (NLS) spanning amino acids 159-169 (GKQPAKKRKAA; score = 11), as well as a second overlapping NLS spanning amino acids 162-170 (PAKKRKAANG; score = 9). In addition, analysis with the NoD predictor identified two putative nucleolar localization (NoLS) sequences [49] spanning amino acids 93-118 and 132-152, suggesting that MoPARP1 contains conserved sequence features associated with nuclear and potential nucleolar targeting (Fig. 1A; Supplementary Table S1).

**Figure 1.**
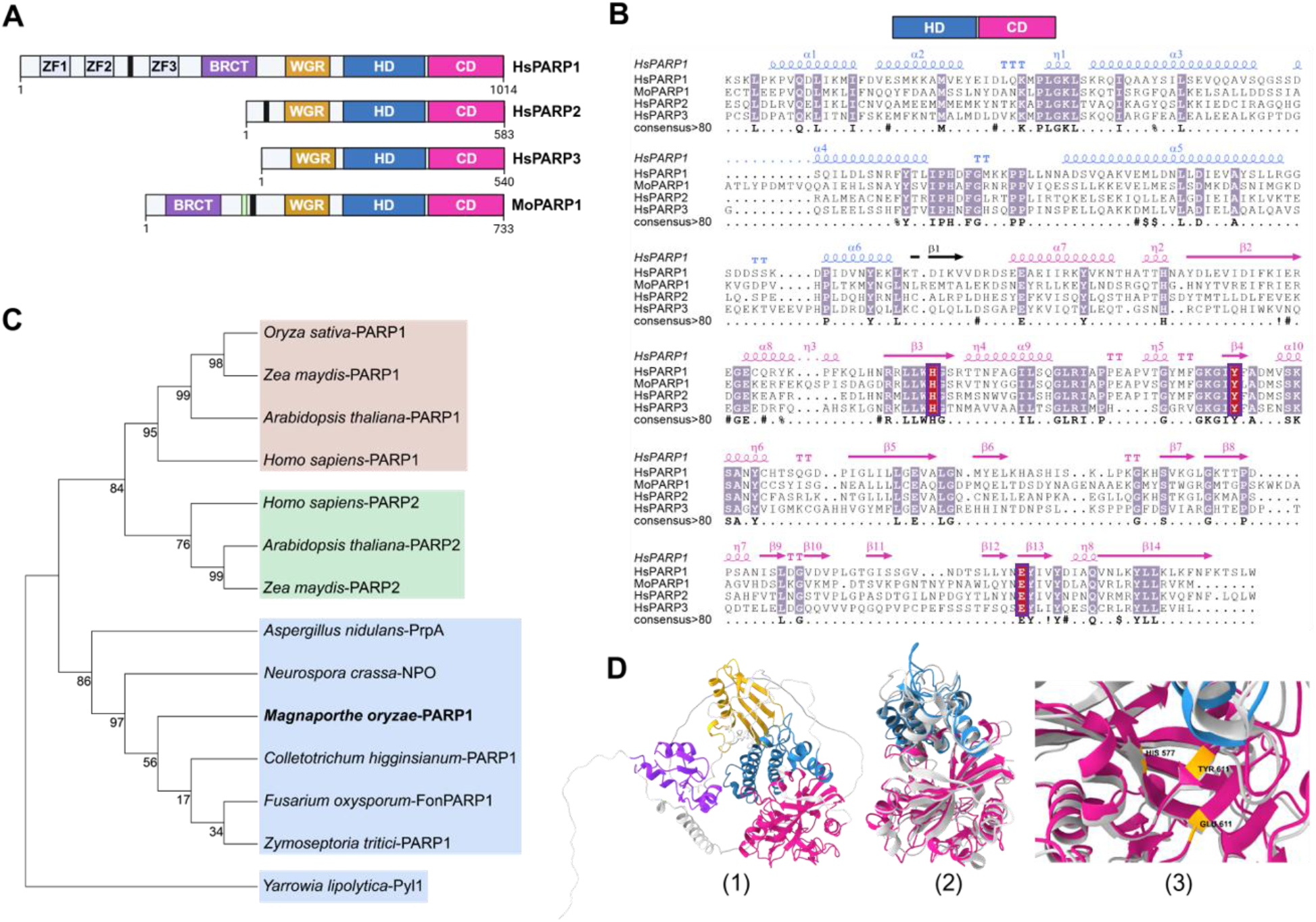
Comparative domain architecture, sequence conservation, phylogenetic relationships, and structural analysis of MoPARP1. **(A)** Schematic representation of the domain architecture of human PARP1 (HsPARP1), PARP2 (HsPARP2), PARP3 (HsPARP3), and *M. oryzae* PARP1 (MoPARP1). Domains are indicated as follows: ZF1-3, zinc finger domains (gray); BRCT, BRCA1 C-terminal domain (purple); WGR, tryptophan-glycine-arginine domain (gold); HD, helical domain (blue); and CD, catalytic domain (magenta). Black boxes indicate predicted nuclear localization signals (NLS), and the small green box indicates a predicted nucleolar localization signal (NoLS). The schematic was generated using BioRender.com. **(B)** Multiple sequence alignment of the HD-CD regions of HsPARP1, HsPARP2, HsPARP3, and MoPARP1. Conserved catalytic H-Y-E residues are highlighted by red boxes. Sequences were aligned using MultAlin [34], and the alignment was visualized using ESPript 3.2 [35] with secondary-structure annotations derived from the HsPARP1 structure (PDB: 7KK2). **(C)** Maximum-likelihood phylogenetic tree of representative PARP homologs from mammals, plants, and fungi based on the conserved HD-CD regions. The tree was generated in MEGA12 [38] using the WAG + F substitution model with a discrete gamma distribution and 1,000 bootstrap replicates. Bootstrap support values are shown at the nodes. **(D)** Structural analysis of MoPARP1. (1) AlphaFold-predicted full-length MoPARP1 structure colored by domain as in panel A. (2) Structural superposition of the predicted MoPARP1 HD–CD region with the experimentally determined HsPARP1 structure (PDB: 7KK2), shown in light gray. (3) Enlarged view of the catalytic site showing conservation of the H–Y–E catalytic residues, including MoPARP1 H577, Y611, and E714. Structural superposition was performed using the Matchmaker tool in UCSF ChimeraX [42,44].

Multiple sequence alignment of the HD-CD regions of MoPARP1 and HsPARP1–3 revealed strong conservation of residues surrounding the catalytic core, including the catalytic H-Y-E motif (Fig. 1B). In MoPARP1, residues H577, Y611, and E714 aligned with the corresponding catalytic residues of the HsPARPs, supporting conservation of the catalytic machinery despite differences in domain architecture.

Because MoPARP1 exhibits PARylation activity [32], phylogenetic analysis focused on representative PARP family members from filamentous fungi, yeasts, plants, and humans with experimentally characterized or well-supported PARylation activity. HsPARP3 was excluded because it primarily catalyzes MARylation rather than PARylation. Maximum-likelihood analysis placed MoPARP1 within the fungal PARP clade, distinct from plant and mammalian PARPs (Fig. 1C). These findings support the classification of MoPARP1 as a fungal member of the PARylating PARP family.

To assess structural conservation, an AlphaFold-predicted model of MoPARP1 was obtained and compared with the experimentally determined structure of the HsPARP1 HD-CD region (PDB: 7KK2) [40,43]. Structural superposition revealed extensive conservation of the catalytic fold and the spatial arrangement of the catalytic H-Y-E residues (Fig. 1D). Structural alignment using the MatchMaker tool in UCSF ChimeraX identified 242 structurally equivalent residues with a root-mean-square deviation (RMSD) of 0.939 Å, indicating a high degree of structural similarity despite substantial sequence divergence [41,42,44]. Notably, MoPARP1 residues H577, Y611, and E714 occupied positions analogous to the catalytic H-Y-E residues of HsPARP1, supporting conservation of the catalytic architecture and catalytic mechanism between fungal and mammalian PARPs.

### E714 is essential for the catalytic activity of MoPARP1

The catalytic glutamate residue within the conserved H-Y-E motif, E714A, is required for ADP-ribosyl transfer in characterized PARP enzymes. To determine whether E714 contributes to MoPARP1 catalytic activity, we generated a site-directed mutant in which glutamate 714 was replaced with alanine (E714A) (Supplemental Fig. S1A). Both recombinant MoPARP1 WT and MoPARP1 E714A were purified to near homogeneity and exhibited comparable profiles (Supplemental Fig. S1B).

To assess catalytic activity, purified proteins were incubated with NAD⁺ and commercially prepared activated DNA (AcDNA), a DNA substrate containing strand breaks that promote PARP activation, and ADPr levels were evaluated by immunoblotting. Consistent with previous findings, MoPARP1 WT exhibited robust PARylation activity in the presence of DNA and NAD⁺, as indicated by the high-molecular-weight signal detected with antibodies recognizing mono- and poly-ADP-ribose (Fig. 2A). In contrast, MoPARP1 E714A failed to produce a detectable PARylation signal, demonstrating that E714 is required for catalytic activity (Fig. 2B). Although equal protein input was verified by BCA assay, the anti-His signal was weaker and more diffuse in catalytically active samples than in reactions lacking PARylation. This difference may reflect reduced accessibility of the His tag following extensive automodification.

**Figure 2.**
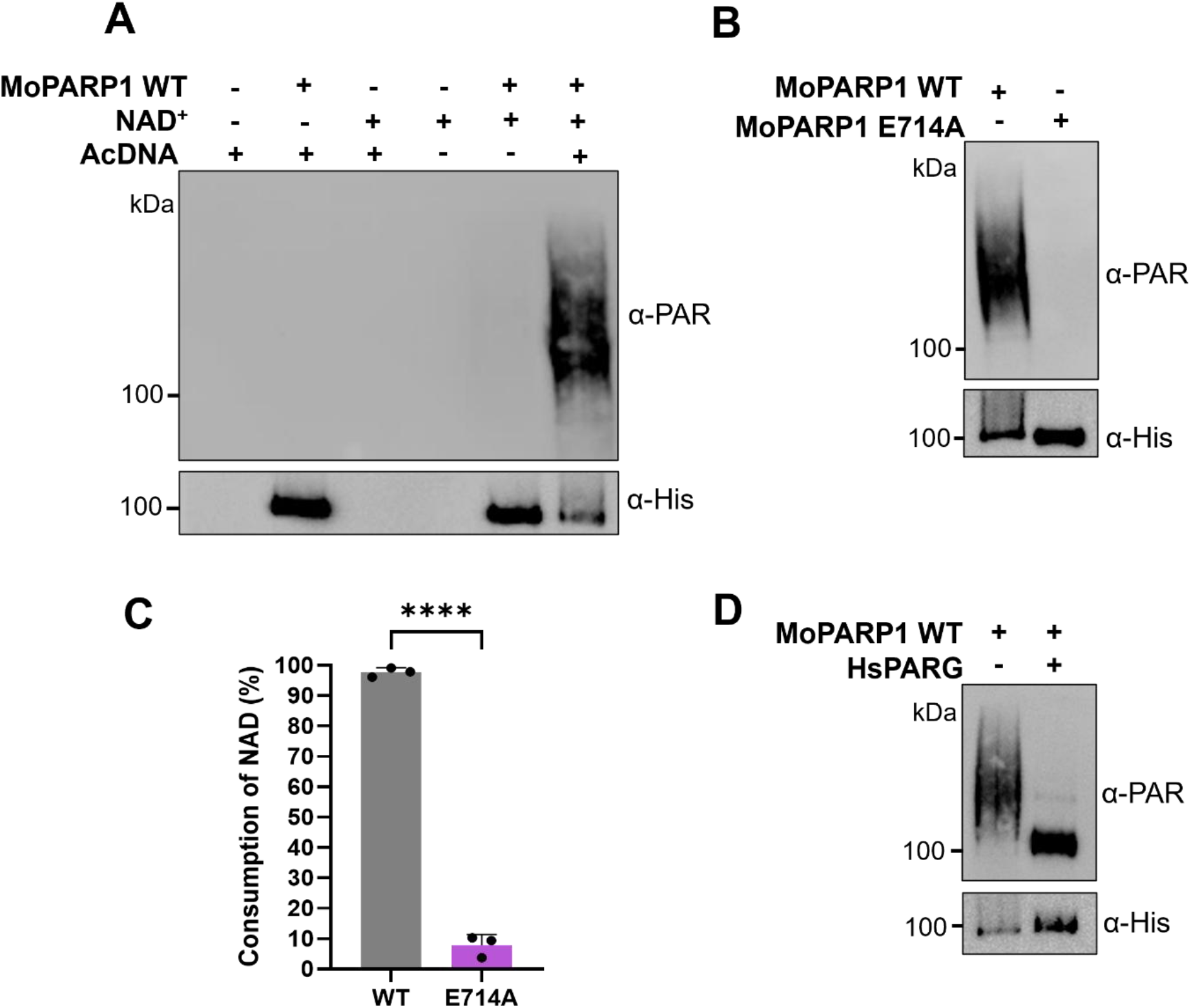
MoPARP1 exhibits DNA-dependent poly(ADP-ribosyl)ation activity in vitro. **(A)** Recombinant MoPARP1 was incubated in the presence or absence of DNA and NAD⁺ to assess DNA-dependent PARylation. **(B)** Comparison of PARylation activity between wild-type (WT) MoPARP1 and the catalytic mutant MoPARP1 E714A. **(C)** NAD⁺ consumption by MoPARP1 WT and MoPARP E714A was measured using a fluorescence-based assay. Bars represent the mean ± SD from three independent experiments (*n* = 3), with individual replicates shown as dots. Statistical significance was determined using a two-tailed unpaired Welch’s *t*-test (****, *P* < 0.0001). **(D)** Removal of MoPARP1-generated poly(ADP-ribose) chains by human poly(ADP-ribose) glycohydrolase (HsPARG). Following a 20-min PARylation reaction, HsPARG was added and incubated for an additional 20 min. For immunoblots, “+” and “−” indicate the presence or absence, respectively, of the indicated reaction components. PARylation was detected with an anti-PAR antibody (α-PAR), and an anti-His antibody (α-His). was used as a loading control.

Because PARylation is coupled with the hydrolysis of NAD^+^, we next quantified catalytic activity of MoPARP1 WT and MoPARP1 E714A by NAD⁺ consumption assay. In the presence of AcDNA, MoPARP1 WT consumed more than 95% of the available NAD^+^, whereas MoPARP1 E714A displayed only basal levels of NAD^+^ consumption (<10%) (Fig. 2C). Together, these results demonstrate that E714 is essential for MoPARP1-mediated NAD⁺ hydrolysis and PARylation activity.

### Hydrolysis of PAR chains is conserved across species

A defining feature of PARylation is its reversibility through the action of PARGs, which hydrolyze PAR chains from modified proteins [50]. Although a functional PARG has not yet been characterized in *M. oryzae*, we sought to determine whether PAR chains synthesized by MoPARP1 could be recognized and hydrolyzed by a heterologous PARG enzyme.

To address this question, PARylated MoPARP1 was incubated with recombinant human PARG (HsPARG) and analyzed by immunoblotting. Treatment with HsPARG resulted in a substantial reduction in PAR signal compared with untreated controls (Fig. 2D). Specifically, the high-molecular-weight PAR smear generated by MoPARP1 WT was substantially diminished following HsPARG treatment, consistent with enzymatic hydrolysis of PAR chains (Fig. 2D). A similar reduction in PAR signal was observed for PARylated HsPARP1 following HsPARG treatment, supporting a comparable pattern of PAR hydrolysis between the fungal and mammalian proteins (Supplementary Fig. S2). These findings demonstrate that PAR synthesized by MoPARP1 is a substrate for HsPARG, indicating that the PAR polymer produced by the fungal enzyme is structurally compatible with the mammalian PAR degradation machinery.

### Human PARP inhibitors suppress MoPARP1 catalytic activity

Previous studies demonstrated that MoPARP1 catalytic activity is inhibited by the HsPARP inhibitor 3-aminobenzamide (3-AB) [32,51]. Consistent with these findings, we observed that 1 and 3 mM 3-AB inhibited more than 70% of MoPARP1 WT PARylation activity (Fig. 3A-C). To determine whether additional HsPARP inhibitors could similarly inhibit MoPARP1 WT, we tested a panel of compounds previously reported to suppress PARP activity in mammalian systems, including olaparib, PJ34, talazoparib, and veliparib [52–56].

**Figure 3.**
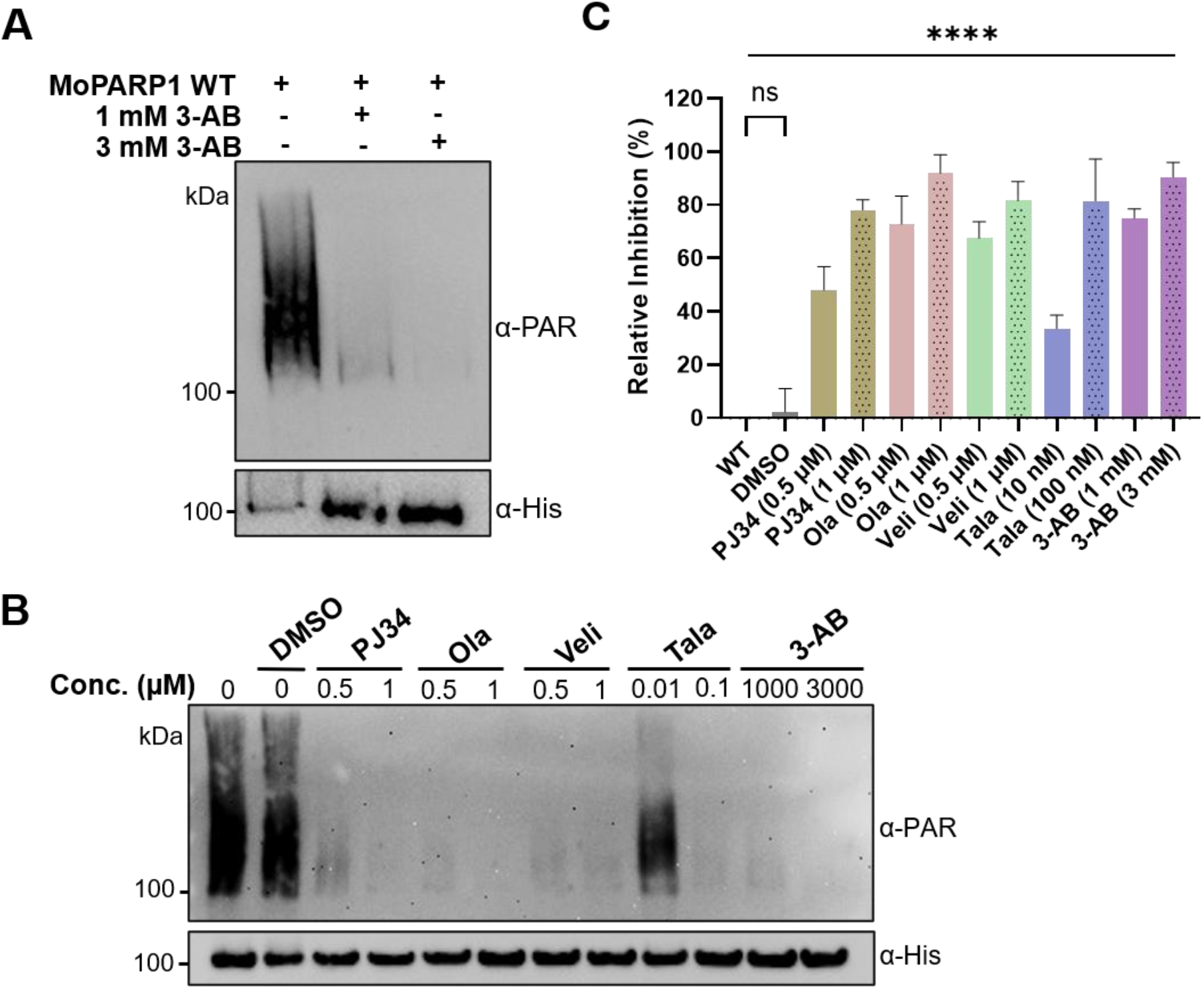
Human PARP inhibitors suppress the catalytic activity of MoPARP1 in vitro. **(A)** Dose-dependent inhibition of MoPARP1 PARylation activity by 3-aminobenzamide (3-AB). Recombinant MoPARP1 was incubated with increasing concentrations of 3-AB in the presence of DNA and NAD⁺, and PARylation was analyzed by immunoblotting using an antibody against poly(ADP-ribose) (α-PAR). Anti-His was used as a protein loading control (α-His). **(B)** Inhibition of MoPARP1 by a panel of mammalian PARP inhibitors. MoPARP1 was incubated with the indicated concentrations of 3-AB, PJ34, olaparib (Ola), veliparib (Veli), and talazoparib (Tala) prior to PARylation reactions. DMSO served as the vehicle control. PARylation was analyzed by immunoblotting using an anti-PAR antibody, and anti-His was used as a protein loading control. **(C)** Quantification of MoPARP1 inhibition following treatment with PARP inhibitors. PAR signal intensities were measured by densitometric analysis of anti-PAR immunoblots using ImageJ. Values were normalized to the DMSO control, which was set to 0% inhibition (100% activity). Data represent the mean ± SD of three independent experiments (*n* = 3). Statistical significance was determined by one-way ANOVA, followed by Dunnett’s multiple comparisons test, with DMSO as the control group. ns, not significant; ****, *P* < 0.0001.

In vitro PARylation assays revealed that multiple mammalian PARP inhibitors suppressed the catalytic activity of MoPARP1 WT, suggesting that key features of the inhibitor-binding pocket are conserved in *M. oryzae* (Fig. 3B-C). Densitometric analysis of PAR signal intensities confirmed that treatment with PJ34, olaparib, veliparib, talazoparib, and 3-AB substantially inhibited MoPARP1 PARylation activity relative to the WT and DMSO controls. Consistent with the immunoblot results, inhibition was observed across all inhibitor classes tested, with several treatments reducing MoPARP1 activity by more than 70%. Notably, talazoparib inhibited MoPARP1 activity at nanomolar concentrations, whereas micromolar or millimolar concentrations were required for the other inhibitors tested (Fig. 3B-C). The DMSO vehicle control did not significantly affect MoPARP1 activity relative to untreated reactions. Collectively, these results demonstrate that MoPARP1 WT is susceptible to multiple structurally distinct mammalian PARP inhibitors and support the conservation of key catalytic features between fungal and mammalian PARPs.

### MoPARP1 recognizes diverse DNA substrates across multiple architectures

An important feature of PARP activity is its dependence on DNA binding, which induces catalytic activation through conformational changes. Our previous NAD⁺ consumption assays confirmed that AcDNA promotes MoPARP1 WT activity (Fig. 2C). We next systematically examined the DNA-binding properties of MoPARP1 WT using a panel of DNA substrates differing in length, strand composition, terminal phosphorylation, and secondary structure. To address these questions, we generated a panel of single-stranded DNA (ssDNA), double-stranded DNA (dsDNA), and structured DNA substrates for electrophoretic mobility shift assays (EMSAs) (Fig. 4A, Supplementary Table S2). DNA-protein complexes were visualized by their reduced migration through agarose gels.

**Figure 4.**
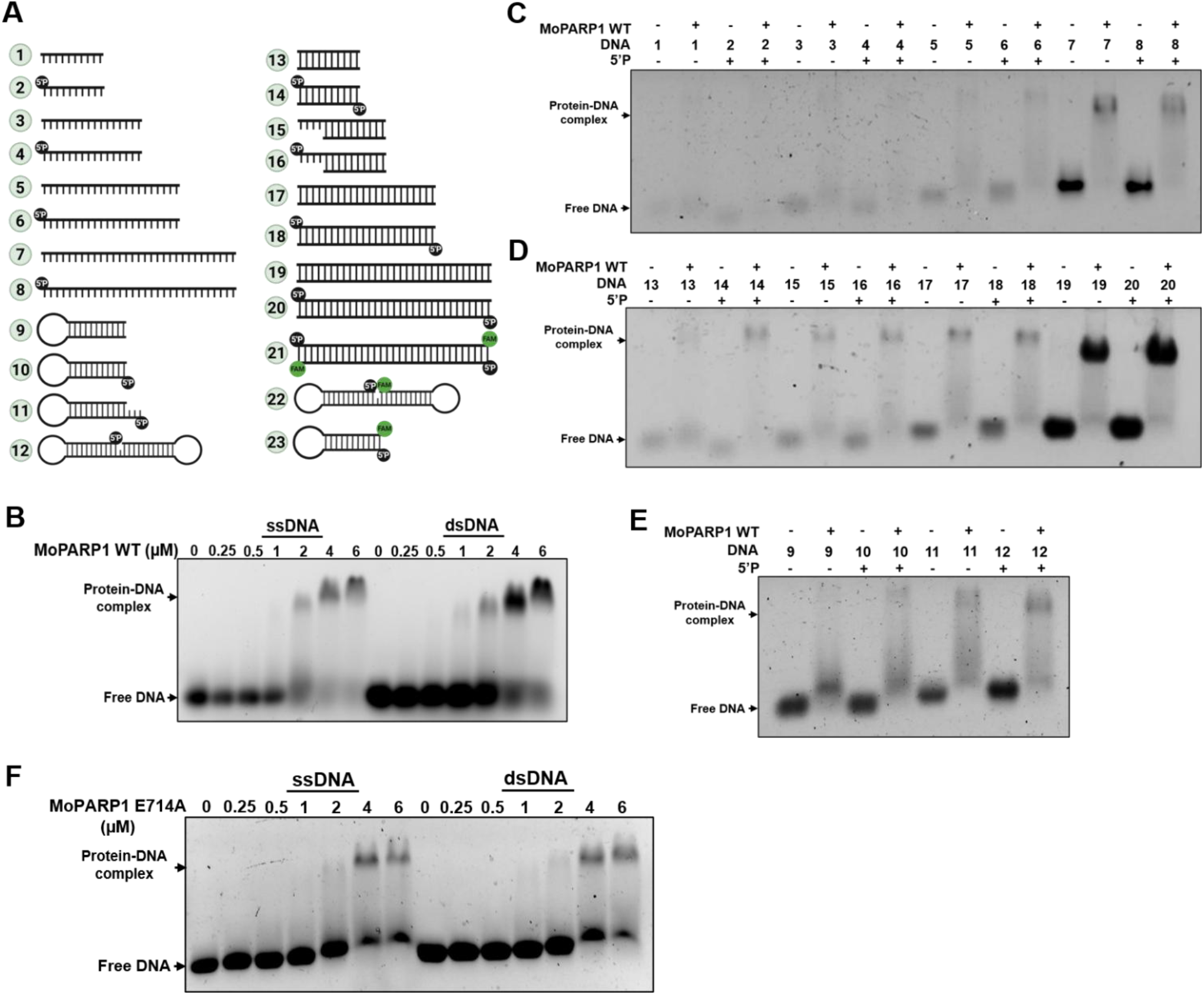
MoPARP1 binds linear and structured DNA substrates. **(A)** Schematic representation of the DNA substrates used in MoPARP1 DNA-binding and catalytic activation assays. 5′P indicates a 5′ phosphate modification, and FAM indicates a 3′ fluorescein (6-FAM) label. **(B)** Agarose electrophoretic mobility shift assay (EMSA) showing concentration-dependent binding of MoPARP1 WT to single-stranded DNA 8 (ssDNA) and double-stranded DNA 20 (dsDNA), while MoPARP1 WT was titrated from 0 to 6 µM. **(C)** Agarose EMSA comparing MoPARP1 WT binding to ssDNA substrates of increasing length with or without a 5′ phosphate modification. **(D)** Agarose EMSA comparing MoPARP1 WT binding to dsDNA substrates of increasing length with or without a 5′ phosphate modification. **(E)** Agarose EMSA comparing MoPARP1 WT binding to structured DNA substrates. **(F)** Agarose EMSA showing concentration-dependent binding of MoPARP1 E714A to ssDNA 8 and dsDNA 22. For panels B-F, DNA substrates were used at 2 µM.

Before comparing binding across the complete substrate panel, we first established an appropriate DNA ratio for the EMSA experiments. The longest linear substrates, the 52-nt ssDNA substrate DNA 8 and the corresponding 52-bp dsDNA substrate DNA 20, were selected for this initial titration. EMSA analysis demonstrated that both substrates formed shifted complexes upon incubation with increasing concentrations of recombinant MoPARP1 WT, consistent with concentration-dependent protein-DNA interactions (Fig. 4B). At DNA ratios of 1:3 or greater, most of the DNA substrate migrated as higher-molecular-weight complexes, indicating robust association of MoPARP1 WT with both ssDNA and dsDNA. Based on these results, a DNA ratio of 1:3, corresponding to 2 µM DNA and 6 µM MoPARP1 WT, was selected for the subsequent substrate comparisons.

To investigate the effects of DNA length and terminal phosphorylation on MoPARP1 WT binding, EMSAs were performed with a panel of ssDNA substrates ranging from 10 to 52 nt in length (Fig. 4C). MoPARP1 WT formed detectable complexes with all ssDNA substrates tested. Longer substrates generally produced more prominently shifted complexes, whereas the presence of a 5′ phosphate had only modest effects on apparent complex formation. Together, these observations suggest that DNA length exerts a greater influence on MoPARP1 association with ssDNA than terminal phosphorylation under the conditions tested. MoPARP1 WT also formed readily detectable complexes with all dsDNA substrates examined (Fig. 4D). Binding to the shortest substrate (10 bp) was comparatively limited, whereas longer substrates produced an increased proportion of higher-molecular-weight complexes. Similar to the ssDNA substrates, 5′ phosphorylation had only modest effects on apparent DNA binding. These results indicate that substrate length is a stronger determinant of DNA association than terminal phosphorylation for both ssDNA and dsDNA substrates.

Having established that MoPARP1 WT binds both ssDNA and dsDNA across a range of lengths, we next sought to determine whether DNA architecture influences substrate recognition. EMSAs were performed using a panel of structured DNA substrates, including hairpin, hanging-end hairpin, and nicked dumbbell conformations (Fig. 4A, E). These substrates were designed to mimic non-canonical DNA structures that arise during DNA repair and replication to evaluate the impact of DNA secondary structure and terminal features on the formation of the MoPARP1-DNA complex. While MoPARP1 WT produced detectable mobility shifts with all structured DNA substrates tested, the extent and migration patterns of the resulting complexes varied among substrates (Fig. 4E). Specifically, the most prominent shifted species were observed with the 18-nt hanging-end hairpin (DNA 11) and with the 38-nt dumbbell DNA substrate (DNA 12). Together, these findings indicate that MoPARP1 WT can recognize diverse structured DNA conformations and suggest that DNA architecture influences the nature of MoPARP1-DNA complexes formed. Notably, MoPARP1 associated with all DNA architectures tested, demonstrating a broad capacity for DNA recognition despite substantial differences in substrate structure.

### Loss of catalytic activity does not abolish MoPARP1 DNA binding

We demonstrated earlier that MoPARP1 E714A retains no detectable PARylation activity, with only baseline NAD⁺ consumption observed (Fig. 2B, C), confirming that substitution of the catalytic glutamate severely disrupts MoPARP1 enzymatic activity. We therefore examined whether the loss of catalysis was accompanied by an impairment in DNA binding. Agarose EMSA analysis showed that MoPARP1 E714A retained the ability to form concentration-dependent complexes with both ssDNA and dsDNA substrates (Fig. 4F). As the concentration of MoPARP1 E714A increased, a greater proportion of each DNA substrate migrated as higher-molecular-weight complexes, indicating continued association of the mutant protein with DNA. Together, these findings demonstrate that MoPARP1 retains DNA-binding capacity in the absence of detectable catalytic activity, indicating that DNA association and catalytic activation represent separable biochemical properties of the enzyme.

### 5’-phosphorylated and structured DNA substrates enhance MoPARP1 catalytic activity

Following our EMSA analyses demonstrating broad DNA-binding behavior across diverse substrates, we sought to determine whether differences in DNA architecture also influence the catalytic activity of MoPARP1 WT. Oligonucleotides previously evaluated in EMSA assays (Fig. 4A) were used for in vitro PARylation assays with recombinant MoPARP1 WT. Equal loading was assessed using BSA rather than using α-his immunoblotting, as α-his detection produced extensive smearing and inconsistent signal intensity, particularly in the samples exhibiting robust catalytic activity.

When incubated with MoPARP1 WT, ssDNA of varying lengths and DNA substrates with secondary structure were able to promote PARylation activity in vitro (Fig. 5A). Interestingly, despite displaying similar binding patterns, our PARylation assays show that DNA substrates of increasing lengths, along with 5’ phosphorylation, enhance MoPARP1 WT PARylation activity. Of note, the smallest DNA substrate tested, DNA 2, produced minimal activation, compared to the largest DNA substrate tested, DNA 8. A similar pattern was observed with the corresponding dsDNA substrates (Fig. 5B).

**Figure 5.**
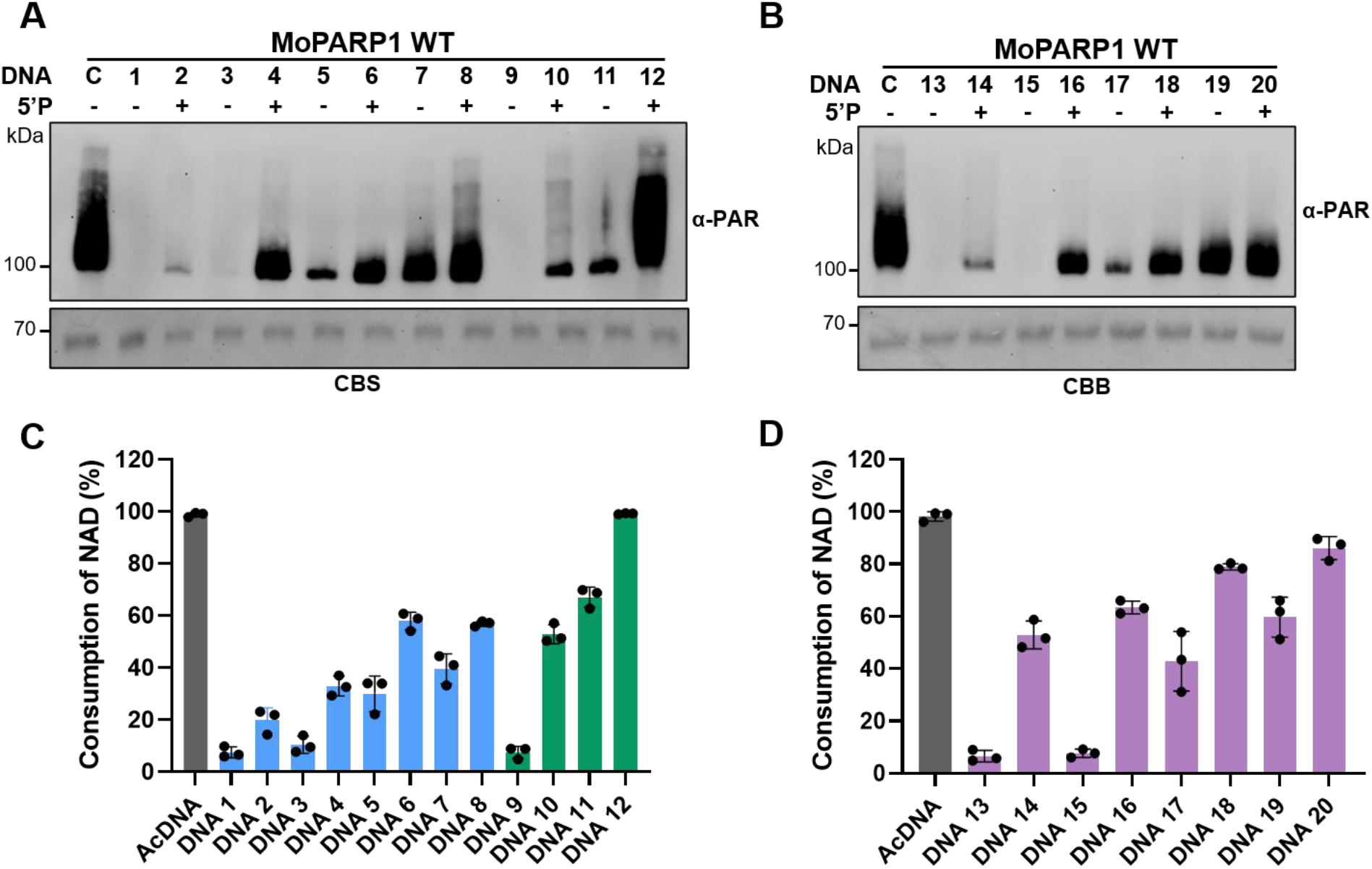
DNA substrates differentially stimulate MoPARP1 catalytic activity. **(A)** In vitro PARylation assay using ssDNA substrates and structured DNA substrates previously evaluated for MoPARP1 WT binding. **(B)** In vitro PARylation assay using dsDNA. For panels A and B, recombinant MoPARP1 WT was incubated with the indicated DNA substrates in the presence of NAD⁺, and PARylation was detected by immunoblotting with an anti-PAR antibody. BSA was included as a loading control and visualized by Coomassie Brilliant Blue (CBB) staining. C denotes a positive control of activated DNA. DNA substrates were used at 1 µM**. (C)** DNA-dependent catalytic activity of MoPARP1 measured by NAD⁺ consumption using the ssDNA substrates shown in panel A. **(D)** DNA-dependent catalytic activity of MoPARP1 measured by NAD⁺ consumption using the double-stranded and structured DNA substrates shown in panel B. For panels C and D, MoPARP1 WT and DNA substrates were used at final concentrations of 150 and 500 nM, respectively. Data are presented as the mean ± SD from three independent biological replicates, with each dot representing one biological replicate. AcDNA denotes activated DNA.

DNA architecture also influenced MoPARP1 WT activity. The 5’-phosphorylated hairpin substrate DNA 10 exhibited more prominent PARylation than its non-phosphorylated counterpart, DNA 9. Moreover, when a hanging-end is introduced to the 5′-phosphorylated hairpin substrate, DNA 11, it further enhanced MoPARP1 WT activity. Among the structured substrates tested, the 5′-phosphorylated nicked dumbbell substrate, DNA 12, produced the most prominent PARylation signal, reaching levels comparable to those observed with AcDNA. Consistent with the in vitro PARylation assays, NAD⁺ consumption measurements showed similar activation patterns by the ssDNA, dsDNA, and structured DNA substrates (Fig. 5C, D). Notably, DNA 12 supported catalytic activity comparable to AcDNA, reaching nearly complete NAD⁺ consumption after 60 min. All substrates containing a 5′ phosphate promoted greater NAD⁺ consumption than their nonphosphorylated substrates of the same length. Together, these results indicate that DNA length, 5′ phosphorylation, and secondary structure influence the catalytic activation of MoPARP1 WT.

### WGR-containing regions support *M. oryzae* PARP1 DNA association

Given that MoPARP1 lacks the canonical N-terminal ZF domains responsible for DNA damage recognition in mammals and that catalytically inactive MoPARP1 retained DNA-binding activity, we sought to determine which regions of MoPARP1 contribute to DNA binding. To address this question, we generated truncated protein constructs representing different regions of MoPARP1, expressed and purified the corresponding recombinant proteins, and assessed their ability to bind DNA by EMSA (Fig. 6A; Supplemental Fig. S3A). The constructs tested included full-length MoPARP1 WT (FL); the N-terminal region containing the BRCT domain (BRCT); the N-terminal BRCT and WGR domains (BRCT-WGR); the PARP domain alone (PARP); the WGR and PARP domains (WGR-PARP); and the WGR domain alone (WGR).

**Figure 6.**
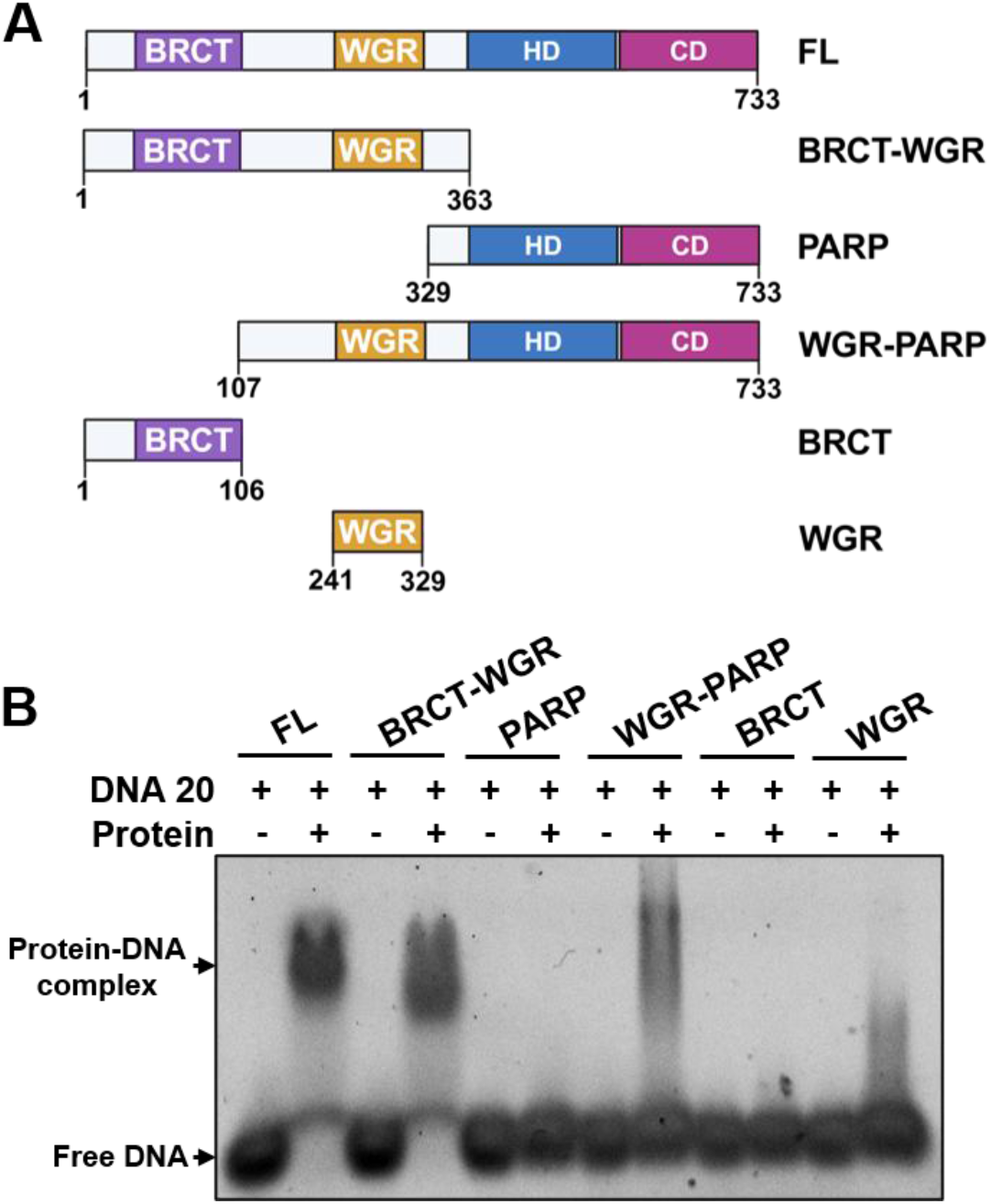
MoPARP1 truncations exhibit differential DNA-binding activity. **(A)** Schematic representation of the MoPARP1 truncation constructs generated and tested in this study. Domains are color-coded as in Fig. 1. The schematic was generated using BioRender.com. **(B)** Agarose EMSA comparing DNA binding by the MoPARP1 truncation constructs shown in panel A. DNA 20 was maintained at 2 µM, and each protein construct was tested at 6 µM.

In the EMSA assays, the BRCT-WGR, WGR-PARP, and WGR constructs produced detectable shifted DNA-protein complexes (Fig. 6B). Specifically, the BRCT-WGR construct produced a compact shifted species, suggesting the formation of a relatively uniform DNA-protein complex. In contrast, the WGR-PARP construct produced a broad vertical smear in complex with DNA, consistent with heterogeneous or unstable complex formation. Similarly, the WGR construct produced the weakest band shift. In contrast, the N-terminal BRCT and the PARP domains alone did not produce detectable shifts under conditions tested. Comparable binding patterns were observed when the BRCT-WGR and WGR-PARP constructs were evaluated using an expanded panel of linear and structured DNA substrates (Supplementary Fig. S3B, C). The stronger and more defined complexes produced by BRCT-WGR compared with WGR alone suggest that N-terminal regions contribute to the stability of DNA-bound complexes. Conversely, the absence of detectable binding by the isolated PARP domain indicates that the catalytic region alone is insufficient for stable DNA association under the conditions tested (Fig. 6B).

### MoPARP1 domain architecture influences DNA-binding and activity

To quantify how MoPARP1 domain composition influences DNA binding, we measured the binding affinities of MoPARP1 WT FL, the catalytically inactive mutant MoPARP1 E714A, and the BRCT-WGR and WGR-PARP truncations. Based on the limited DNA binding observed in the agarose EMSA, the WGR truncation was excluded from further quantitative binding analysis. Proteins were tested against three DNA substrates: DNA 21, a blunt-ended DNA; DNA 22, a nicked dumbbell substrate; and DNA 23, a hairpin substrate. These substrates correspond to the FAM-labeled derivatives of DNA 20, DNA 12, and DNA 10, respectively. These substrates were selected because they represent distinct DNA damage or end configurations that PARP proteins may encounter during DNA repair [57,58]. Apparent dissociation constants (*K_d_*) were determined by native PAGE-EMSA assays using 3’-FAM-labeled DNA oligonucleotide substrates and incubated with increasing protein concentrations (Supplementary Fig. S4). Here, we show that MoPARP1 WT FL bound all three DNA structures with nanomolar affinity, with apparent *K_d_* values of approximately 43-47 nM (Fig. 7A, B; Supplementary Fig. S5A). The MoPARP1 E714A mutant displayed comparable affinities across all substrates (44-50 nM), further indicating that DNA binding does not depend on catalytic activity (Fig. 7A, B, S5B).

**Figure 7.**
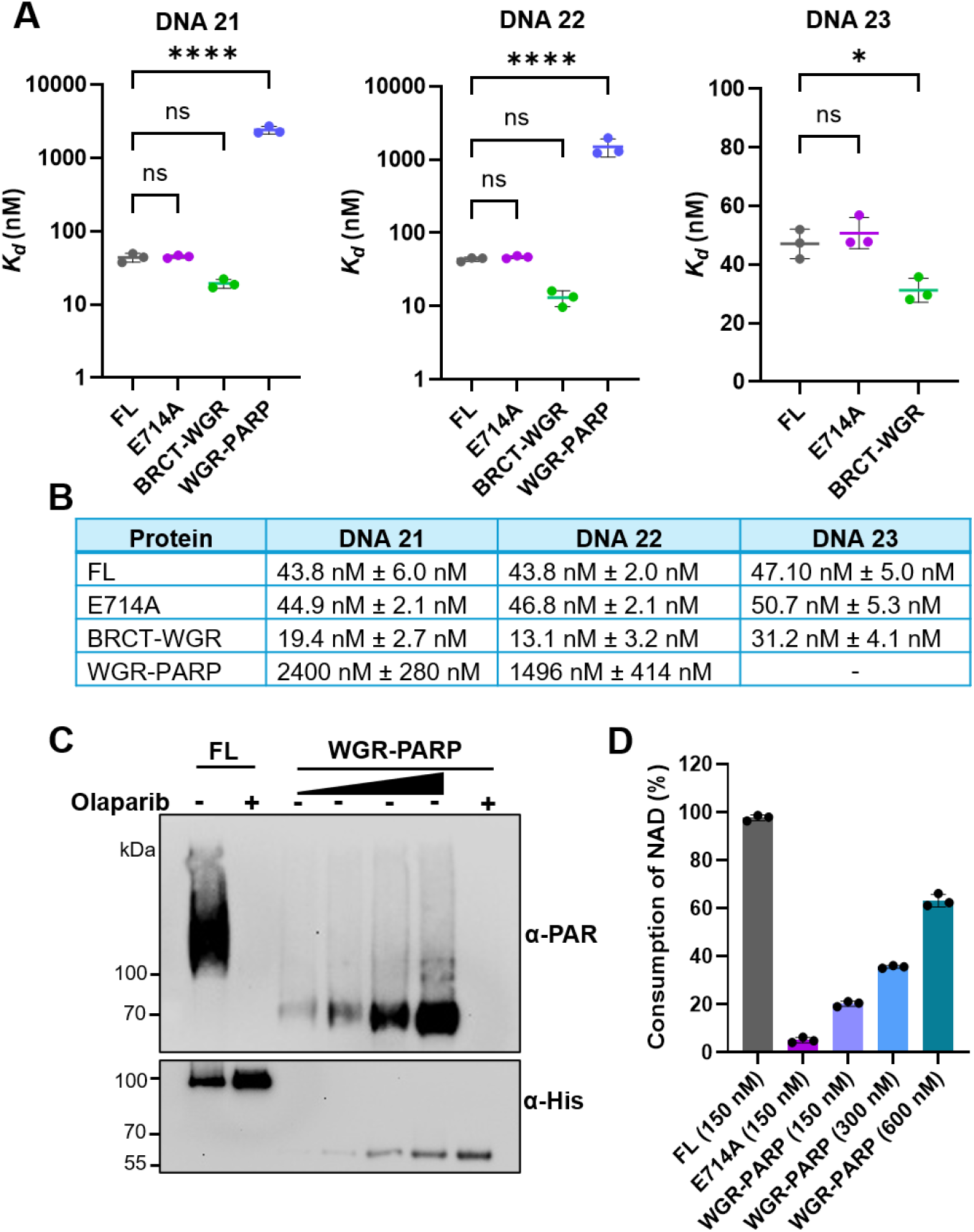
DNA-binding affinity and catalytic activity of MoPARP1 constructs. **(A)** Dissociation constants (*K*d) were determined for full-length MoPARP1 WT (FL), the catalytically inactive E714A mutant, BRCT-WGR, and WGR-PARP binding to three different DNA substrates. *K*d values are presented as the mean ± SD from three independent biological experiments. Statistical significance was determined by one-way ANOVA followed by Dunnett’s multiple-comparisons test, with FL used as the control. Significance is indicated as ns, not significant; *P* < 0.05 (*); and *P <* 0.0001 (****). **(B)** Summary table showing the mean *K*d ± SD for each protein construct and DNA substrate (*n* = 3 biological replicates). –, *Kd* not determined. **(C)** In vitro PARylation activity of FL and WGR-PARP proteins. The WGR-PARP construct was tested at concentrations ranging from 100 to 1000 nM. Activated DNA was used as a DNA substrate. Where indicated, reactions contained 1 µM olaparib. PARylation was detected with an anti-PAR antibody, and an anti-His antibody was used as a loading control. **(D)** Measurement of NAD⁺ consumption by FL, E714A, and WGR-PARP in the presence of activated DNA. FL and E714A was used at 150 nM, while WGR-PARP was titrated from 150-600 nM.

Native PAGE EMSA analysis of MoPARP1 truncations revealed that apparent DNA binding was strongly influenced by domain architecture. The BRCT-WGR construct showed higher affinity relative to MoPARP1 FL for all three DNA substrates, with apparent *K_d_* values of approximately 19.4 nM for blunt-ended (DNA 21), 13.1 nM for nicked dumbbell (DNA 22), and 31.2 nM for the hairpin substrate (DNA 23) (Fig. 7A, B; Supplementary Fig. S5C). In contrast, the WGR-PARP construct displayed markedly reduced affinity for blunt-ended (DNA 21) and nicked dumbbell DNA (DNA 22), with apparent *K_d_* values in the micromolar range, while an apparent *K_d_* for the hairpin substrate (DNA 23) could not be determined under the conditions tested (Fig. 7A, B; Supplementary Fig. S5D). Together, these findings indicate that the WGR domain contributes to MoPARP1 DNA binding, whereas high-affinity DNA association requires additional N-terminal elements beyond the minimal WGR-PARP core.

Next, to determine which domains are necessary for catalytic activity, MoPARP1 constructs were evaluated using an in vitro PARylation assay with AcDNA. As previously shown, MoPARP1 WT exhibited robust ADPr activity, whereas the catalytically inactive MoPARP1 E714A mutant had no detectable signal (Fig. 2C, Supplementary Fig. S3D). The BRCT-WGR construct also yielded no detectable signal, consistent with the absence of the catalytic domain (Fig. 7C, Supplementary Fig. S3D). Interestingly, the PARP construct also yielded no detectable signal, suggesting that the catalytic domain alone is insufficient to support detectable PARylation under the conditions tested. In contrast, the MoPARP1 WGR-PARP construct retained in vitro catalytic activity, demonstrating that the WGR and catalytic domain are sufficient to promote DNA-dependent PARylation. Increasing the concentration of WGR-PARP protein resulted in progressively greater PARylation, consistent with a concentration-dependent increase in catalytic activity (Fig. 7C, Supplementary Fig. S3D). Additionally, olaparib treatment strongly inhibited PARylation activity in both MoPARP1 FL and WGR-PARP constructs, indicating that the activity of both proteins remains sensitive to PARP inhibition.

To further quantify catalytic activity, NAD⁺ consumption was measured (Fig. 7D). Similar to our previous findings, MoPARP1 WT showed over 90% consumption of NAD⁺, whereas the MoPARP1 E714A mutant showed minimal consumption. Consistent with the immunoblotting results, the WGR-PARP construct exhibited concentration-dependent NAD⁺ depletion as protein concentration increased from 150 to 600 nM. Together, these findings indicate that the WGR domain, with the catalytic domain, is sufficient to support DNA-dependent PARylation.

### DNA-binding affinity does not predict MoPARP1 catalytic activation

To determine whether DNA-binding affinity directly predicts the magnitude of MoPARP1 catalytic activation, we compared the catalytic properties of three DNA substrates that exhibited similar apparent binding affinities in quantitative EMSA assays (Fig. 7B; Supplementary Fig. S5A). MoPARP1 bound DNA 21, DNA 22, and DNA 23, corresponding to the FAM-labeled derivatives of DNA 20, DNA 12, and DNA 10, respectively, with comparable apparent *K_d_* values of approximately 43-47 nM, indicating little difference in equilibrium DNA association among these substrates. Despite these similar binding affinities, the corresponding unlabeled substrates produced markedly different NAD⁺ consumption profiles (Fig. 8A). After 60 min, DNA 12 and our positive control, AcDNA, supported nearly complete NAD⁺ consumption (∼99% and ∼98%, respectively), whereas DNA 20 and DNA 10 reached approximately 85% and 52% NAD⁺ consumption, respectively.

**Figure 8.**
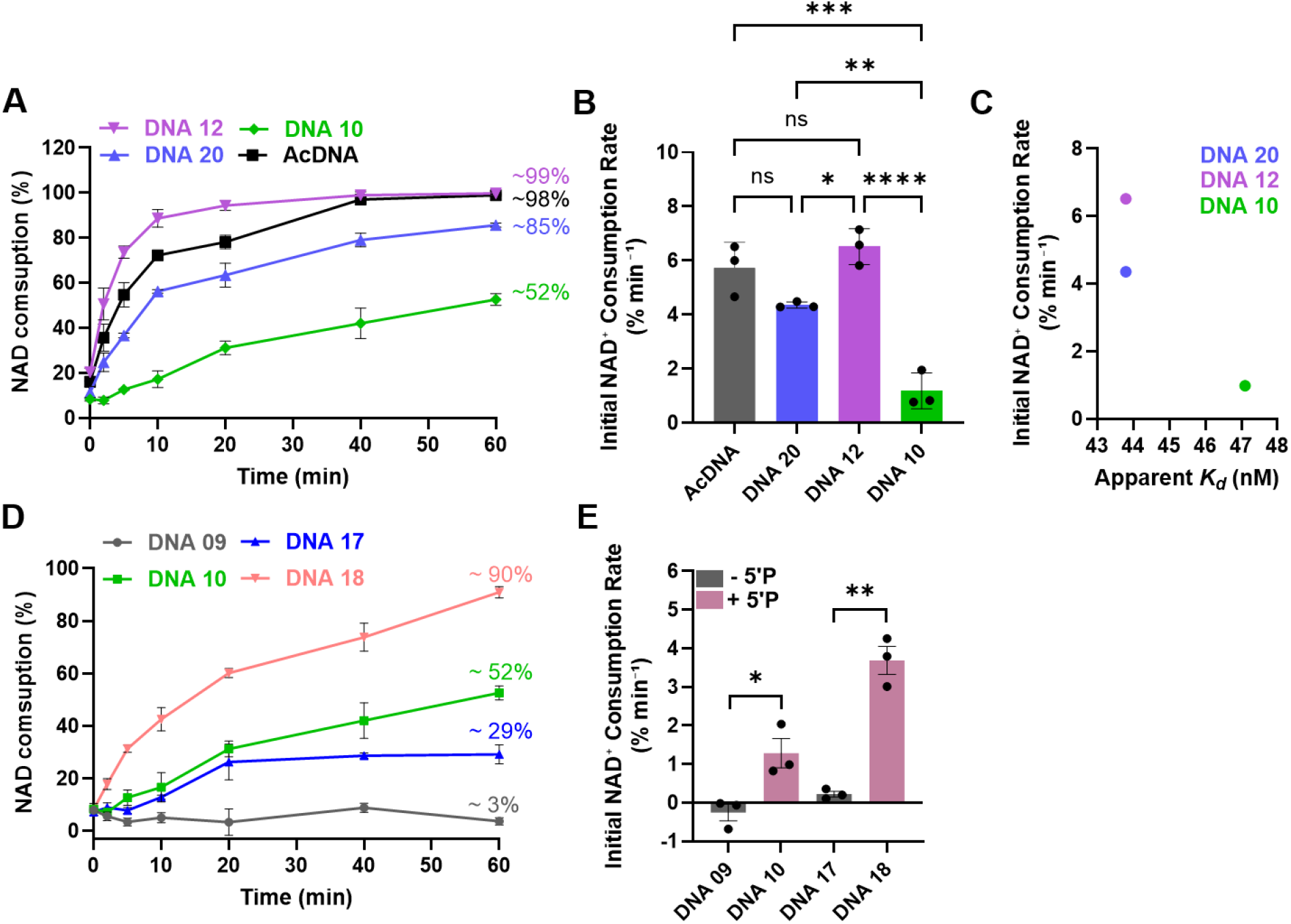
DNA-binding affinity does not predict MoPARP1 catalytic activation. **(A)** Time-dependent NAD⁺ consumption by MoPARP1 WT in the presence of AcDNA, DNA 12, DNA 20, and DNA 10. NAD⁺ consumption was monitored over time and expressed as the percentage of NAD⁺ consumed relative to the starting concentration. **(B)** Initial NAD⁺ consumption rates (*v₀*) were determined by linear regression of the initial linear phase of the reactions. **(C)** Relationship between apparent *Kd* and catalytic activation. Mean apparent *Kd* values were plotted against mean initial NAD⁺ consumption rates (*v₀*) for DNA 10, DNA 20, and DNA 12. Despite exhibiting similar apparent DNA-binding affinities (∼44-47 nM), these substrates produced markedly different catalytic outputs, indicating that equilibrium DNA-binding affinity alone does not predict catalytic activation. **(D)** Time-dependent NAD⁺ consumption by MoPARP1 WT in the presence of DNA 9, DNA 10, DNA 17, and DNA 18. DNA 10 and DNA 18 contain a 5′ phosphate, whereas DNA 9 and DNA 17 represent the corresponding non-phosphorylated substrates. NAD⁺ consumption was monitored over time and expressed as the percentage of NAD⁺ consumed relative to the starting concentration. **(E)** Initial NAD⁺ consumption rates (*v₀*) were determined by linear regression of the initial linear phase of the reactions for DNA 9, DNA 10, DNA 17, and DNA 18. Data represents mean ± SD from three independent experiments (*n* = 3). Statistical significance in (B) was determined by one-way ANOVA followed by Tukey’s multiple-comparisons test. Statistical significance in (E) was determined by an unpaired Welch’s *t*-test. ns, not significant; *P* < 0.05; *P* < 0.01; *P* < 0.001; *P* < 0.0001.

To quantitatively compare catalytic output during the initial phase of the reaction, initial NAD⁺ consumption rates (*v₀*), expressed as % NAD⁺ consumed min⁻¹, were determined by linear regression of the first 10 min of the reaction (Fig. 8B). DNA 12 exhibited the highest initial consumption rate (*v₀* = 6.51% NAD⁺ consumed min⁻¹), followed by DNA 20 (*v₀* = 4.35% NAD⁺ consumed min⁻¹), whereas DNA 10 promoted substantially lower activity (v₀ = 1.18% NAD⁺ consumed min⁻¹). Plotting apparent *K_d_* values against *v₀* further highlighted the disconnect between DNA binding and catalytic activation (Fig. 8C).

Despite exhibiting similar apparent DNA-binding affinities, DNA 10, DNA 20, and DNA 12 produced markedly different catalytic outputs. Together, these findings demonstrate that equilibrium DNA-binding affinity alone is insufficient to predict productive catalytic activation by MoPARP1.

### 5′ phosphorylation enhances MoPARP1 catalytic activation in a DNA-context-dependent manner

To further examine the contribution of 5′ phosphorylation to MoPARP1 activation, we compared NAD⁺ consumption kinetics using matched DNA substrate pairs that differed only in the presence or absence of a 5′ phosphate (Fig. 8D, E). In this case, DNA 9 and DNA 17 are unphosphorylated, while DNA 10 and DNA 18 contain a 5’ phosphate. For both substrate pairs, phosphorylated DNA promoted greater NAD⁺ consumption throughout the reaction time course than the corresponding non-phosphorylated substrate (Fig. 8D). After 60 min, DNA 10 and DNA 18 supported approximately 52% and 90% NAD⁺ consumption, respectively, whereas the corresponding non-phosphorylated substrates, DNA 9 and DNA 17, reached only approximately 3% and 29% NAD⁺ consumption. These results indicate that 5′ phosphorylation enhances MoPARP1 catalytic activity across distinct DNA architectures.

To quantify catalytic activity during the initial phase of the reaction, initial NAD⁺ consumption rates (v₀) were determined by linear regression of the first 10 min of the reaction (Fig. 8E). DNA 9 exhibited little or no detectable catalytic activity, whereas the corresponding 5′-phosphorylated substrate DNA 10 supported an initial NAD⁺ consumption rate of approximately 1.20% NAD⁺ consumed min⁻¹. Similarly, the initial consumption rate increased from approximately 0.22% NAD⁺ consumed min⁻¹ for DNA 17 to 3.68% NAD⁺ consumed min⁻¹ for DNA 18. In both cases, the presence of a 5′ phosphate significantly enhanced MoPARP1 catalytic activation. However, the magnitude of stimulation differed substantially between the two DNA substrate pairs, indicating that the effects of 5′ phosphorylation are influenced by the underlying DNA architecture. Together, these findings demonstrate that 5′ phosphorylation promotes MoPARP1 activation but is not sufficient on its own to determine catalytic output, supporting roles for both terminal chemistry and DNA structural context in regulating enzymatic activity.

## Discussion

PARPs are highly conserved enzymes that coordinate cellular responses to DNA damage through the synthesis of PAR chains using NAD⁺ as a substrate in a process known as PARylation [8,59,60]. While mammalian PARPs have been extensively characterized, comparatively little is known regarding the biochemical properties and biological functions of PARPs in filamentous fungi, particularly in plant pathogens. Here, we provide a detailed biochemical characterization of MoPARP1 from the filamentous fungal plant pathogen *Magnaporthe oryzae,* supported by comparative sequence and structural analyses. Our findings demonstrate that MoPARP1 is a bona fide DNA-dependent PARP that retains conserved catalytic features while displaying distinct domain requirements for DNA recognition and catalytic activation. Importantly, our results indicate that stable DNA association and productive catalytic activation are mechanistically distinguishable properties of MoPARP1. Collectively, these results establish a mechanistic framework for understanding PARylation in fungal pathogens and provide a biochemical foundation for future studies examining how MoPARP1 contributes to genome maintenance and pathogenic fitness.

DNA recognition is central to the activation of DNA-dependent PARPs [11,61,62]. Our DNA-binding experiments showed that MoPARP1 WT interacts with a wide range of DNA substrates, including ssDNA, dsDNA, and structured DNA. The binding to structured DNA, such as hairpins or nicks, suggests that MoPARP1 may respond to strand interruptions or unusual DNA conformations that can arise during DNA replication and repair [11,62,63]. These preferences resemble the damage-associated mechanisms described in mammalian DNA-dependent PARPs, suggesting MoPARP1 may play a role in responding to a variety of damaged DNA structures during genome maintenance. Thus, although MoPARP1 lacks the ZF domains of HsPARP1, it retains comparatively broad DNA-binding capacity.

Our results also indicate that DNA binding and catalytic activation are not equivalent properties of MoPARP1. The presence of a 5′ phosphate produced only modest differences in apparent DNA binding, whereas 5′ phosphorylation and DNA architecture had substantially greater effects on catalytic activity. Moreover, structurally distinct DNA substrates exhibited comparable apparent binding affinities yet produced markedly different levels of NAD⁺ consumption. These observations indicate that DNA-binding affinity alone does not determine the magnitude of MoPARP1 catalytic activation. Instead, DNA architecture and terminal chemistry appear to influence the efficiency of a DNA-bound MoPARP1 complex in reaching a catalytically competent state. This distinction is consistent with DNA-dependent allosteric activation mechanisms described for mammalian PARPs [61,64].

This distinction becomes particularly apparent when MoPARP1 is compared with other DNA-dependent PARPs. Mammalian HsPARP2/3 exhibit strong preferences for specific 5′-phosphorylated DNA breaks, with WGR-mediated interactions contributing to both lesion recognition and catalytic activation [11,45,62,65]. Similarly, recent characterization of PARP1 from the human pathogen *Aspergillus fumigatus* (Af-PARP1) demonstrated preferential activation of selected 5′-phosphorylated nicked, gapped, and overhanging DNA substrates, indicating that DNA structure-dependent activation is also a feature of fungal PARPs [33]. Our findings extend this emerging model by showing that MoPARP1 associates with a broader range of DNA architectures, including duplex DNA, while catalytic activation remains strongly influenced by DNA architecture and terminal chemistry. Thus, conservation of fungal PARP domain architecture does not necessarily impose a uniform DNA-recognition profile. Although phosphorylated substrates generally promoted greater NAD⁺ consumption than their non-phosphorylated counterparts, the extent of activation remained highly dependent on DNA context. Furthermore, DNA substrates with similar apparent binding affinities generated markedly different catalytic outputs, indicating that productive activation cannot be explained solely by DNA association. The closer phylogenetic relationship of MoPARP1 to PARPs from other fungal pathogens, such as those from *Colletotrichum higginsianum* and *Neurospora crassa,* further supports the existence of a distinct fungal lineage of PARP enzymes that may have diversified in regulatory mechanisms and substrate specificity [66,67]. Together with recent reports describing functional PARPs in *A*. *fumigatus*, *Fusarium oxysporum*, and *Yarrowia lipolytica*, these findings suggest that fungal PARPs retain conserved catalytic cores while exhibiting substantial diversification in DNA recognition and regulatory properties.

The domain architecture underlying DNA recognition varies substantially among members of the PARP family. HsPARP1 recognizes DNA strand breaks primarily through its N-terminal ZF domains, whereas HsPARP2/3 lacks ZF domains and largely relies on WGR-mediated interactions to bind nicked, gapped, and dsDNA [11,67]. Sequence alignment and domain architecture analysis revealed that MoPARP1 contains the conserved BRCT, WGR, regulatory helical, and catalytic domains characteristic of DNA-associated PARPs [61,68,69]. However, MoPARP1 lacks the canonical N-terminal ZF domains present in HsPARP1 [11,68]. This suggested that MoPARP1 may recognize DNA primarily through WGR-mediated interactions, similar to the DNA recognition mechanisms utilized by HsPARP2/3 [11,65]. Indeed, domain truncation analysis revealed the WGR domain as a central determinant of detectable MoPARP1 DNA binding. All constructs that produced detectable mobility shifts contained the WGR domain, whereas constructs lacking WGR failed to produce detectable shifts under the conditions tested, including the isolated catalytic region. These observations support a central contribution of WGR to MoPARP1 DNA recognition, although they do not exclude additional contributions from neighboring domains in the context of the full-length protein.

In mammalian HsPARP2/3, WGR also contributes to coupling DNA engagement to catalytic activation, suggesting that the MoPARP1 WGR region may perform a similar dual function [68]. Upon DNA binding, PARPs undergo DNA-dependent interdomain rearrangements that alter regulatory elements within the catalytic region, facilitating access of NAD⁺ to the active site and stimulating PAR synthesis and downstream PARylation events involved in chromatin remodeling and DNA repair [61,68,70]. HsPARP1, for example, is rapidly activated following recognition of DNA strand breaks and subsequent remodeling of regulatory elements within the catalytic domain [61,68,70]. These studies establish interdomain communication and DNA-dependent allosteric activation as central features of mammalian PARP regulation and provide a framework for how the DNA architecture of MoPARP1 may couple DNA recognition to catalytic activation [61,68,69,71]. These studies establish interdomain communication and DNA-dependent allosteric activation as central features of mammalian PARP regulation. Sequence alignment and domain architecture analysis revealed that MoPARP1 contains the conserved BRCT, WGR, regulatory helical, and catalytic domains characteristic of DNA-associated PARPs [61,68,69]. While sharing this core functional architecture with HsPARP1/2/3, MoPARP1 lacks the canonical N-terminal ZF domains present in HsPARP1 [11,68]. This suggested that MoPARP1 may recognize DNA primarily through the WGR domain, similar to the DNA recognition mechanisms utilized by HsPARP2 and HsPARP3.

Of particular interest, the BRCT-WGR construct showed the highest apparent DNA binding affinity among the constructs tested. Compared with the weaker binding observed for the WGR and WGR-PARP constructs, this finding suggests that the N-terminal BRCT-containing region may enhance or stabilize WGR-dependent DNA association in MoPARP1. Importantly, the isolated BRCT construct did not produce detectable DNA binding under our assay conditions, indicating that BRCT alone is insufficient for stable MoPARP1 DNA association. Its strong effect in the BRCT-WGR construct, therefore, suggests that the BRCT-containing region enhances DNA association in the context of WGR rather than functioning as an autonomous high-affinity DNA-binding module. This observation is related to, but mechanistically distinct from, recent findings for Af-PARP1, in which the BRCT domain enhances DNA association and contributes to efficient catalytic activation [33]. Thus, the MoPARP1 data are consistent with a cooperative contribution of the N-terminal BRCT-containing region and WGR to DNA binding, while suggesting that the relative contributions of BRCT and WGR have diversified among fungal PARPs. Additional targeted experiments will be required to define the specific role of BRCT.

An additional distinction emerged when the DNA-binding and catalytic properties of the MoPARP1 truncations were considered together. Although BRCT-WGR displayed high-affinity DNA binding, the WGR-PARP construct exhibited substantially weaker apparent DNA affinity yet retained DNA-dependent PARylation activity. In contrast, the catalytic PARP domain alone was unable to support detectable activity. These observations suggest that stable DNA association and catalytic competence are supported by partially distinct domain configurations. Thus, the domain arrangement supporting the highest apparent DNA-binding affinity is not identical to the minimal configuration required for catalytic signaling. Such a dual role for WGR is consistent with its central contribution to DNA-dependent allosteric activation in HsPARP1/2/3 [62,71]. The MoPARP1 WGR-PARP region may therefore represent a minimal DNA-responsive catalytic unit, whereas additional N-terminal regions promote high-affinity DNA association. Further studies will be required to define the precise contributions of individual domains to DNA binding and catalytic activation.

In addition to the conserved PARP-associated domains, MoPARP1 contains predicted nuclear and nucleolar localization signals. Although their functionality was not experimentally validated in the present study, these motifs are consistent with the expected localization of proteins involved in genome maintenance and DNA repair. Future studies examining MoPARP1 localization during growth, stress responses, and plant infection may provide additional insight into the cellular contexts in which fungal PARylation operate.

Consistent with prior reports, MoPARP1 displayed robust DNA-dependent PARylation activity in vitro [32,51]. Structural comparisons between MoPARP1 and HsPARP1 revealed extensive conservation within the catalytic core, including preservation of the canonical H-Y-E catalytic triad responsible for NAD⁺ cleavage and PAR synthesis [71]. Specifically, these comparisons identified H577, Y611, and E714 as the corresponding H-Y-E motif residues in MoPARP1. Mutation of the conserved catalytic glutamate within the catalytic pocket, E714, of MoPARP1 completely abolished detectable PARylation and strongly reduced NAD^+^ consumption, confirming that this residue is essential for MoPARP1 catalytic activity. Importantly, DNA binding was retained in the catalytically inactive MoPARP1 E714A mutant, demonstrating that catalytic activity is not required for DNA engagement. The retention of DNA binding by E714A further supports the conclusion that DNA association and catalytic activation represent separable steps in MoPARP1 function.

This separation of DNA recognition from activation has parallels in other DNA-dependent PARPs. WGR-mediated interactions in HsPARP2 and HsPARP3 contribute both to lesion recognition and communication with the catalytic region [11]. Likewise, mutation of a conserved WGR tyrosine in Af-PARP1 abolishes 5′-phosphate-dependent activation while preserving DNA binding [33]. In MoPARP1, separation between DNA association and allosteric activation is evident both at the protein level through E714A and at the substrate level through DNA structures with similar apparent affinity but different catalytic output.

Although a canonical *M. oryzae* PARG homolog has not yet been functionally characterized, PAR synthesized by MoPARP1 was efficiently hydrolyzed by HsPARG, indicating that fungal PAR linkages share sufficient structural conservation with mammalian PAR to be recognized by the mammalian PAR-degrading machinery [7,72]. Together with direct detection of PAR, the susceptibility of MoPARP1-generated products to HsPARG further supports the conclusion that MoPARP1 synthesizes polymeric ADP-ribose rather than exclusively mono-ADP-ribose products. Thus, despite divergence in DNA-recognition and regulatory mechanisms, structural features of the PAR polymer itself appear sufficiently conserved for cross-species PARG recognition. Nevertheless, identification and characterization of endogenous PAR-degrading activities in *M. oryzae* will be important for determining how PAR signal duration and turnover are regulated in the fungal cell.

In addition to conservation in PAR synthesis and hydrolysis, MoPARP1 was sensitive to HsPARP inhibitors, such as 3-AB, PJ34, olaparib, veliparib, and talazoparib [32,51,73]. All inhibitors suppressed MoPARP1 catalytic activity in vitro, supporting conservation of structural features within the catalytic NAD⁺-binding pocket that permit recognition of chemically diverse PARP inhibitors. PARP inhibitors act by occupying the NAD⁺-binding pocket and blocking PAR synthesis [73–75]. Although mammalian PARP inhibitors differ in their PARP-trapping activities [76,77], trapping was not evaluated in this study, and therefore, the effects of these compounds on MoPARP1-DNA complex stability remain unknown. The sensitivity of MoPARP1 to mammalian PARP inhibitors highlights both its conservation and the challenge of selective targeting. Direct exploitation of the highly conserved catalytic pocket could also affect host PARPs. Consequently, fungal-selective inhibition may require identification of structural or allosteric features that distinguish fungal PARPs from their mammalian counterparts. The DNA-recognition architecture of MoPARP1, including its dependence on WGR and the high-affinity binding observed for the BRCT-WGR region, may provide alternative regulatory features for future investigations. The contrast between conservation of the catalytic pocket and divergence of the DNA-recognition machinery suggests that regulatory interfaces involved in DNA sensing or allosteric activation may provide more fungus-selective opportunities than the NAD⁺-binding site itself. Whether these features can be selectively exploited remains to be determined.

Together, these results provide mechanistic insight into MoPARP1 domain function, DNA recognition, catalytic regulation, PAR turnover, and inhibitor sensitivity. MoPARP1 retains a highly conserved catalytic core while displaying distinct requirements for DNA recognition and activation. Importantly, structurally distinct DNA substrates can exhibit comparable binding affinities but markedly different catalytic outputs, suggesting that DNA engagement alone does not determine productive PAR synthesis. Collectively, these findings support a model in which WGR-containing domain architecture mediates DNA engagement, N-terminal regions enhance or stabilize high-affinity DNA association, and DNA architecture together with 5′-terminal chemistry influences the efficiency of productive catalytic activation.

Importantly, all biochemical assays described here were performed with recombinant protein under in vitro conditions, and current structural comparisons rely primarily on computational modeling. Future structural and cellular studies will therefore be needed to determine how the biochemical properties identified here relate to MoPARP1 function in the full-length protein and native fungal context. This work defines key biochemical and regulatory features of MoPARP1 and provides a mechanistic framework for understanding DNA-dependent PARylation in a filamentous fungal plant pathogen. These findings broaden our understanding of genome-maintenance mechanisms in phytopathogenic fungi and identify regulatory features of fungal PARPs that may ultimately be relevant to strategies for controlling fungal plant disease.

## Acknowledgments

We would like to thank the Fernandez laboratory members for their support in reviewing this manuscript and providing their constructive feedback.

## Author Contributions

N.P. (Formal analysis [lead], Investigation [lead], Validation [supporting], Visualization [lead], Writing –original draft [lead]), R.E.K. (Formal analysis [supporting], Investigation [supporting], Writing –review & editing [supporting]), A.R.K. (Investigation [supporting], Writing –review & editing [supporting]), and J.F. (Conceptualization [lead], Formal analysis [supporting], Methodology [lead], Project administration [lead], Resources [lead], Supervision [lead], Validation [lead], Writing –review & editing [lead]).

## Data Availability Statements

The data underlying this article are available in the article and in its online supplementary material.

