## Supplemental Materials for "Biochemical and Mechanistic Characterization of the DNA-dependent Poly(ADP-ribose) Polymerase, MoPARP1, in *Magnaporthe oryzae*"

**Supplemental Data**

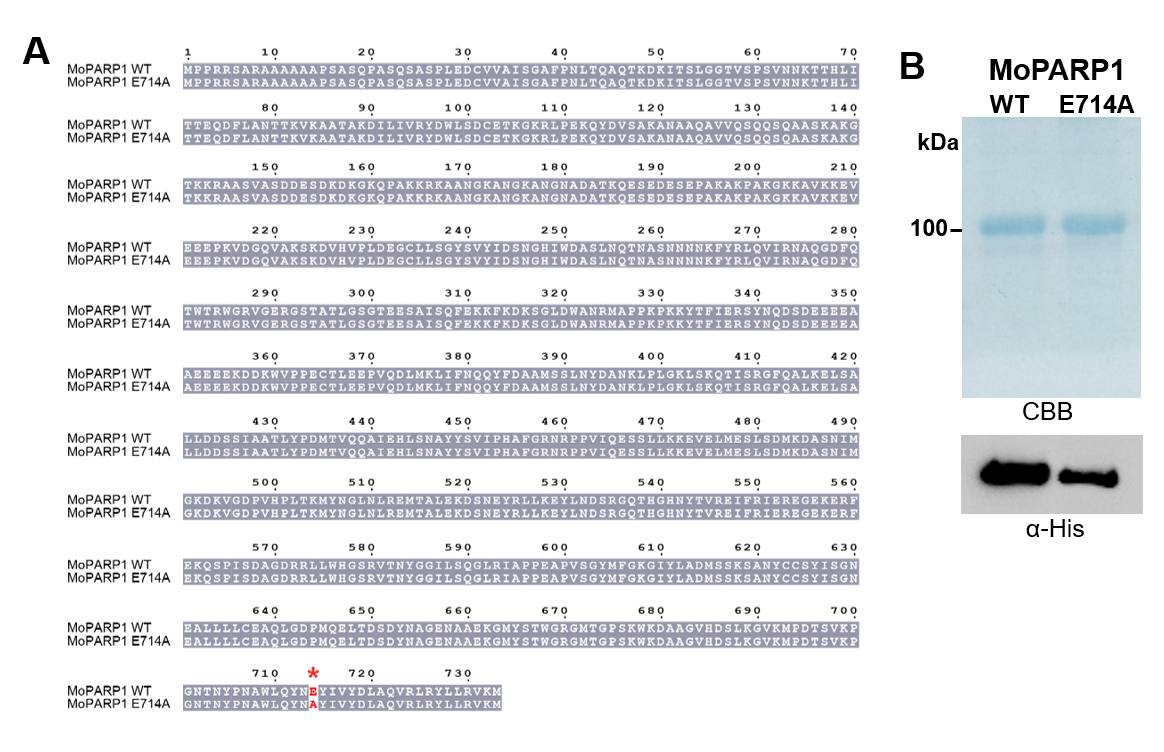

**Supplemental Figure S1. Generation and purification of recombinant MoPARP1 wild-type and catalytic mutant proteins.** **(A)** Sequence alignment of MoPARP1 WT and MoPARP1 E714A. A red asterisk indicates the catalytic glutamic acid residue (E714) replaced by alanine. **(B)** Analysis of purified recombinant MoPARP1 WT and MoPARP1 E714A following nickel-affinity and size-exclusion chromatography. Purified proteins were visualized by Coomassie Brilliant Blue (CBB) staining and detected by immunoblotting with an anti-His antibody (α-His). Protein size for MoPARP1 is 86.5 kDa.

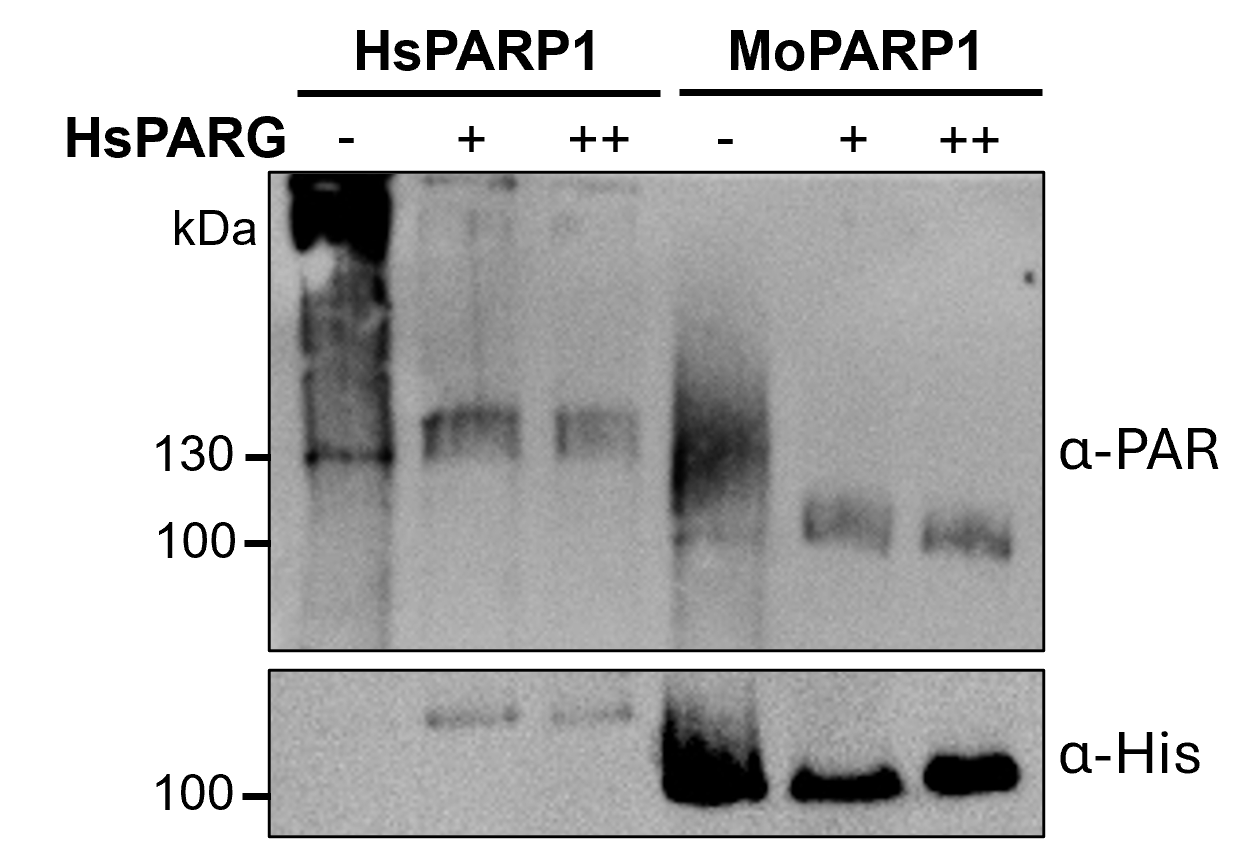

**Supplemental Figure S2. PARG-mediated removal of PAR from HsPARP1 and MoPARP1.** Recombinant HsPARP1 and MoPARP1 were allowed to undergo PARylation for 20 min at room temperature and subsequently untreated (-) or treated with 20 (+) or 50 (++) nM human PARG (HsPARG) for 30 min at 30 °C. PARylation was detected by immunoblotting with an anti-PAR antibody (α-PAR), and protein levels were assessed using an anti-His antibody (α-His). Size of proteins are as follows: HsPARP1: 114.5 kDa, MoPARP1: 86.5 kDa.

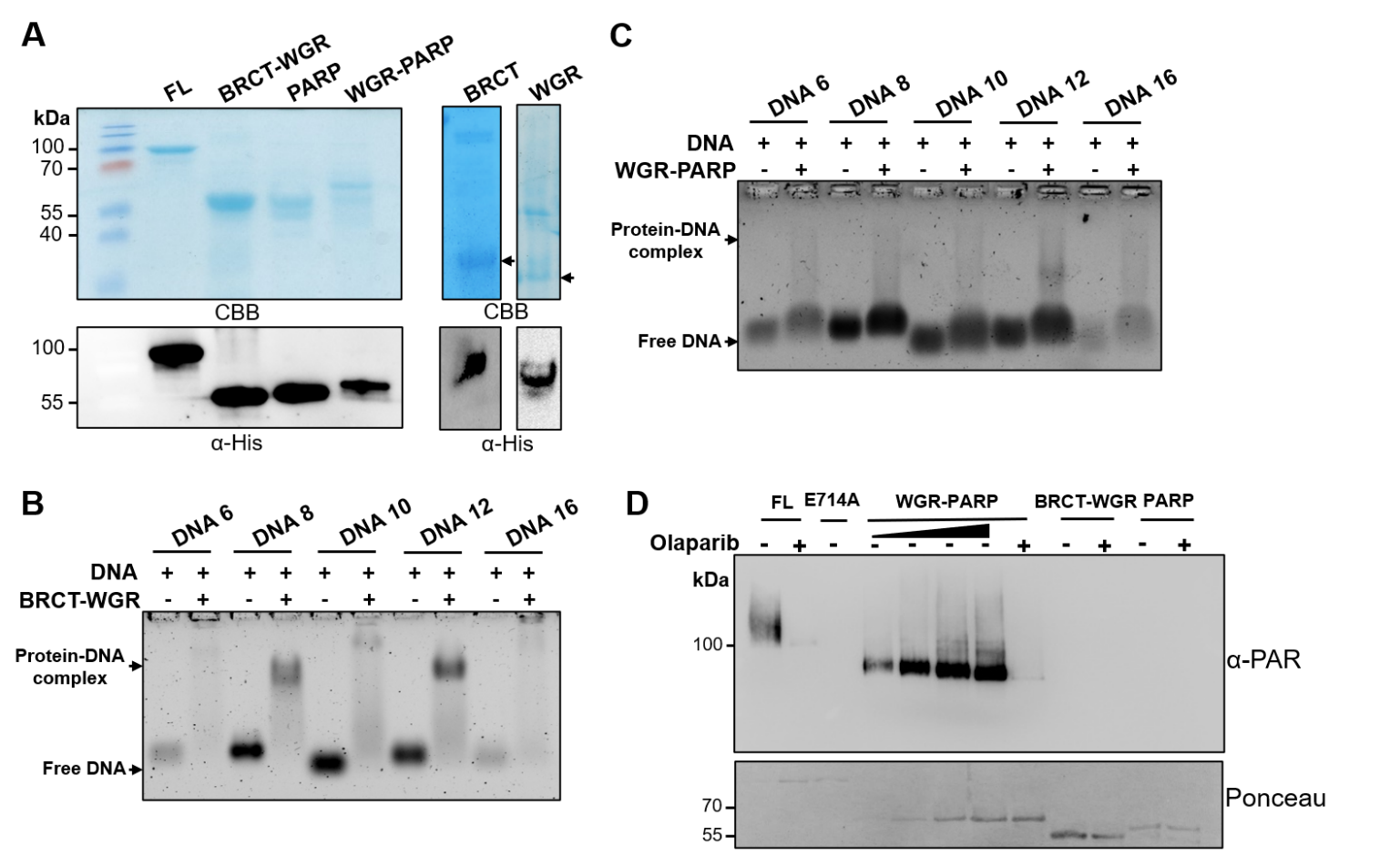

**Supplemental Figure S3. Generation, purification, and DNA-binding analysis of recombinant MoPARP1 truncation constructs. (A)** Purification of recombinant MoPARP1 truncation constructs, including full-length MoPARP1 (FL), BRCT-WGR, PARP, and WGR-PARP, following nickel-affinity and size-exclusion chromatography. Purified proteins were visualized by Coomassie Brilliant Blue (CBB) staining and detected by immunoblotting with an anti-His antibody (α-His). **(B-C)** EMSA analysis of DNA binding by the recombinant BRCT-WGR (B) and WGR-PARP (C) constructs. Proteins were incubated with the indicated DNA substrates (DNA 6, DNA 8, DNA 10, DNA 12, and DNA 16) at a DNA:protein ratio of 1:3 (2 μM DNA and 6 μM protein), and DNA-protein complexes were resolved by agarose gel electrophoresis. **(D)** In vitro PARylation assay using protein truncations, FL, E714A, WGR-PARP, BRCT-WGR, and PARP. Activated DNA was used as a DNA substrate. Where indicated, reactions contained 1 µM olaparib. PARylation was detected with an anti-PAR antibody, and proteins were visualized in the membrane by Ponceau staining. For panel D, 100 nM of FL and E714A were used, whereas WGR-PARP was titrated from 100 to 1000 nM. BRCT-WGR and PARP truncations were used at fixed 1000 nM concentration. Protein truncation sizes are as follows: FL – 86.5 kDa, BRCT-WGR – 43.9 kDa, PARP – 49 kDa, WGR-PARP – 60.5 kDa, BRCT – 15.3 kDa, WGR – 16.2 kDa.

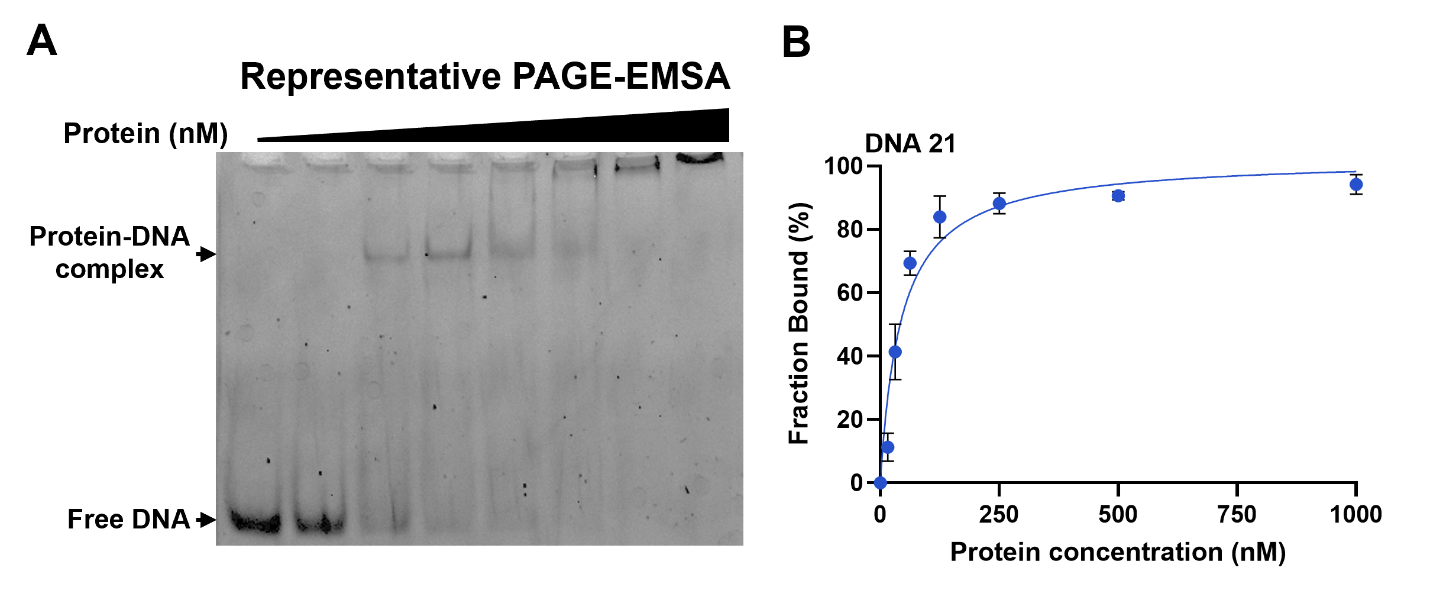

**Supplemental Figure S4. Representative analysis of MoPARP1 DNA-binding affinity by electrophoretic mobility shift assay.** **(A)** Representative native PAGE-EMSA showing binding of increasing concentrations of MoPARP1 WT to DNA 21. DNA 21 was maintained at 5 nM, while MoPARP1 WT was titrated from 0 to 1 nM. Free DNA and protein-DNA complexes are indicated. **(B)** Corresponding binding curve generated from quantification of the fraction of DNA bound at each MoPARP1 concentration. Dissociation constants (*K_d_*) were determined by nonlinear regression using a one-site specific binding model in GraphPad Prism. DNA 21 is shown as a representative example of the *K_d_* analysis. Data are presented as mean ± standard deviation (SD) from three biological replicates.

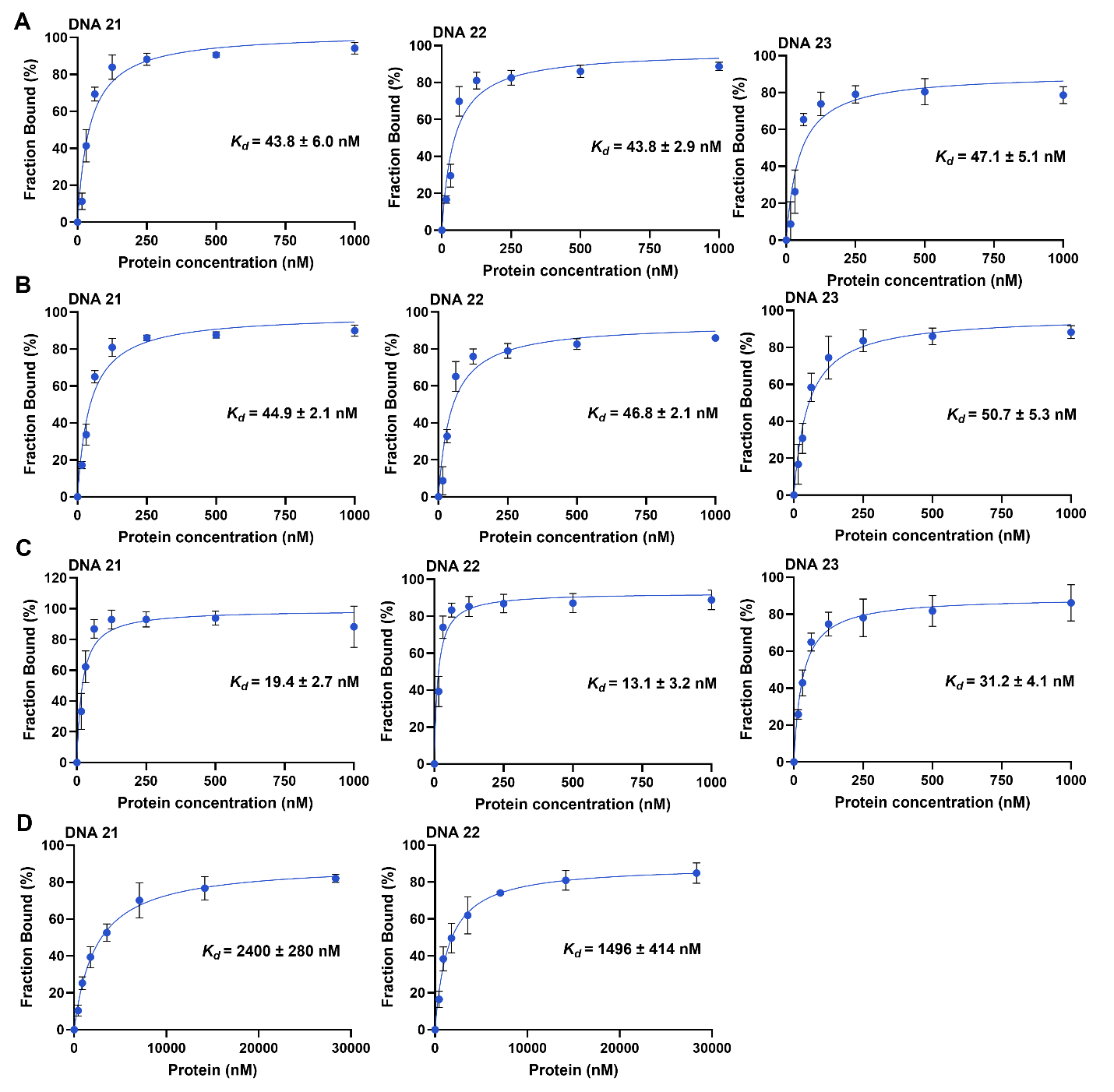

**Supplemental Figure S5. PAGE-EMSA analysis of DNA binding by MoPARP1 proteins.** Representative binding curves are shown for **(A)** WT, **(B)** E714A, **(C)** BRCT-WGR, and **(D)** WGR-PARP. DNA was maintained at a fixed concentration of 5 nM. WT, E714A, and BRCT-WGR were titrated from 1 to 1000 nM, whereas WGR-PARP was titrated from 1 to 30,000 nM. Fraction bound was quantified from native PAGE-EMSA assays and plotted as a function of protein concentration. Binding data were fit by nonlinear regression using a one-site specific binding model in GraphPad Prism. Three independent biological replicates are shown, and dissociation constants (*K_d_*) are reported as mean ± standard deviation (SD; *n* = 3).

**Supplemental Table S1**. Predicted nuclear and nucleolar localization signals in MoPARP1.

| **Feature** | **Prediction Tool** | **Position (aa)** | **Score** |
| --- | --- | --- | --- |
| Monopartite NLS | NLS Mapper | 159–169 | 11 |
| Overlapping NLS | NLS Mapper | 162–170 | 9 |
| Putative NoLS1 | NoD | 93–118 | N/A |
| Putative NoLS2 | NoD | 132–152 | N/A |

**Supplemental Table S2**. Oligonucleotide sequences used in this study.

| **Name** | **Primer Sequence (5'-3')** | **Source** |
| --- | --- | --- |
| DNA 1 | GCCTATAGGC | [1] |
| DNA 2 | (5'P) GCCTATAGGC | [1] |
| DNA 3 | CGGTCGCCTATAGGC | This study |
| DNA 4 | (5'P) CGGTCGCCTATAGGC | This study |
| DNA 5 | GCCACTAGTCTCGCAGTTAGCGCG | This study |
| DNA 6 | (5'P) GCCACTAGTCTCGCAGTTAGCGCG | This study |
| DNA 7 | GCCACTAGTCTCGCAGTTAGCGCGTATACGCGCTAACTGCGAGACTAGTGGC | [1] |
| DNA 8 | (5'P)GCCACTAGTCTCGCAGTTAGCGCGTATACGCGCTAACTGCGAGACTAGTGGC | [1] |
| DNA 9 | GGTAGTTCATTGAACCTACC | This study |
| DNA 10 | (5'P)GGTAGTTCATTGAACCTACC | This study |
| DNA 11 | (5'P) TCAGCGGTAGTTCATTTGAACCTACC | This study |
| DNA 12 | (5'P)GGAAGTTCTTTTGAACTTCCGCGAAGCTTTTGCTTCGA | [1] |
| DNA 13 | Annealed DNA 1 | [1] |
| DNA 14 | Annealed DNA 2 | [1] |
| DNA 15 | Fwd: CGGTCGCCTATAGGC  Rev: GCCTATAGGC | This study |
| DNA 16 | Fwd: (5'P)CGGTCGCCTATAGGC  Rev: GCCTATAGGC | This study |
| DNA 17 | Annealed DNA 5 | This study |
| DNA 18 | Annealed DNA 6 | This study |
| DNA 19 | Annealed DNA 7 | [1] |
| DNA 20 | Annealed DNA 8 | [1] |
| DNA 21 | Annealed DNA 14 | [1] |
| DNA 22 | DNA 12 with 3'FAM | [1] |
| DNA 23 | DNA 10 with 3'FAM | This study |
